# Insulin receptor activation by proinsulin preserves synapses and vision in retinitis pigmentosa

**DOI:** 10.1101/2020.05.13.092833

**Authors:** Alonso Sánchez-Cruz, Alberto Hernández-Pinto, Concepción Lillo, Carolina Isiegas, Miguel Marchena, Ignacio Lizasoain, Fátima Bosch, Pedro de la Villa, Catalina Hernández-Sánchez, Enrique J. de la Rosa

## Abstract

Synaptic loss, neuronal death, and circuit remodeling are common features of central nervous system neurodegenerative disorders. Retinitis pigmentosa (RP), the leading cause of inherited blindness, is a group of retinal dystrophies characterized by photoreceptor dysfunction and death. The insulin receptor, a key controller of metabolism, also regulates neuronal survival and synaptic formation, maintenance, and activity. Indeed, deficient insulin receptor signaling has been implicated in several brain neurodegenerative pathologies. We present evidence linking impaired insulin receptor signaling with RP. We describe a selective decrease in the levels of the insulin receptor and its downstream effector phospho-S6 in retinal horizontal cell axons in the rd10 mouse model of RP, as well as aberrant synapses between rod photoreceptors and the postsynaptic terminals of horizontal and bipolar cells. A gene therapy strategy to induce sustained proinsulin production restored retinal insulin receptor signaling, by increasing S6 phosphorylation, without peripheral metabolic consequences. Moreover, proinsulin preserved photoreceptor synaptic connectivity and prolonged visual function in electroretinogram and optomotor tests. These findings support the therapeutic potential of proinsulin in RP.

## INTRODUCTION

Neurodegenerative disorders are complex pathological conditions that involve, among other processes, synaptic loss and neuronal cell death leading to deterioration of neuronal structure and function. According to the 2017 Global Burden of Disease report (Collaborators, 2019), neurodegenerative diseases are the second leading cause of death and a major cause of disability. The development of strategies to cure or at least delay the progression of neurodegenerative diseases has been hindered by their diverse etiology and complex nature, and by the limited regenerative capacity of neurons. There is thus an urgent need for effective medical interventions. As part of the central nervous system (CNS), the retina shares multiple pathophysiological features with the brain (de La Rosa & Hernandez-Sanchez, 2018). Despite their diverse etiology, retinal neurodegenerative diseases, like those affecting the brain, are characterized by synaptic failure and neuronal cell death (Pfeiffer, Marc et al., 2020). Retinitis pigmentosa (RP) comprises a group of hereditary retinal neurodegenerative conditions with a complex genetic etiology. To date more than 60 genes and 3,000 mutations have been implicated in RP, and over 300 genes associated with inherited retinal dystrophies (https://sph.uth.edu/retnet/disease.htm) have been identified. RP is characterized by primary dysfunction and death of photoreceptor cells followed by reactive gliosis and remodeling of the retinal structure, resulting in vision loss and eventual blindness (Cuenca, Fernandez-Sanchez et al., 2014, Pfeiffer et al., 2020). RP is categorized as a rare disease (prevalence 1/3,500-4,000), but accounts for most cases of hereditary blindness. Gene therapy would be the ideal definitive treatment, and one such therapy has been recently approved for a related retinal dystrophy (Russell, Bennett et al., 2017). However, the complexity and diversity of the mutations underlying RP necessitate the development of alternative therapeutic approaches, particularly those independent of the causative mutation, as well as palliative treatments. The insulin receptor (INSR), traditionally considered a key peripheral metabolic regulator, is increasingly viewed as an important modulator of neuronal cell survival, and synaptic formation, maintenance, and activity (Arnold, Arvanitakis et al., 2018, Banks, Owen et al., 2012, Chiu & Cline, 2010). Since initial reports described widespread INSR expression in the CNS, including the retina (de la Rosa, Bondy et al., 1994, Havrankova, Roth et al., 1978, Marks, Porte et al., 1990, Rodrigues, Waldbillig et al., 1988, Unger, McNeill et al., 1989), INSR signaling has been implicated in a growing number of CNS functions. In addition to regulating feeding behavior and peripheral metabolism, central INSR signaling is involved in memory formation and cognitive functions (Arnold et al., 2018, Banks et al., 2012, Gralle, 2017). The mechanisms underpinning the non-metabolic actions of INSR are being gradually unraveled. At the neuronal level, INSR is involved in the control of synaptic function, through regulation of neurotransmitter receptor trafficking, and in synapse maintenance and dendritic arbor formation (Chiu & Cline, 2010, Lee, Huang et al., 2011). Downregulation of INSR signaling has been implicated in several neurodegenerative diseases (Arnold et al., 2018), particularly Alzheimer’s disease (Moloney, Griffin et al., 2010, Rivera, Goldin et al., 2005, Steen, Terry et al., 2005, Talbot, Wang et al., 2012), and INSR stimulation proposed as a potential treatment for neurodegenerative disorders (Chapman, Schioth et al., 2018).

We previously showed that INSR stimulation in the embryonic retina promotes neuronal differentiation and downregulates developmental cell death (Hernandez-Sanchez, Lopez-Carranza et al., 1995, Valenciano, Corrochano et al., 2006), and that the insulin precursor proinsulin (Pi) can slow RP progression (Corrochano, Barhoum et al., 2008, Fernandez-Sanchez, Lax et al., 2012, Isiegas, Marinich-Madzarevich et al., 2016). In the present study, we investigated the role of INSR in retinal neurodegeneration and sought to characterize the neuroprotective role of Pi as a putative INSR ligand. We present the first evidence linking impaired INSR signaling with retinal neurodegeneration, and provide insight into the multifaceted neuroprotective role of INSR. We employed a gene therapy strategy to produce sustained increases in systemic Pi levels without peripheral metabolic consequences. Pi treatment restored INSR signaling as measured by S6 phosphorylation, preserved photoreceptor synaptic connectivity, and more importantly, extended visual function in a murine model of RP.

## RESULTS

### Insulin receptor expression and signaling in WT and rd10 retinas

The insulin receptor gene (*Insr*) is expressed in mammals as two differentially spliced mRNA isoforms that differ by the presence of a small exon (exon 11) encoding 12 amino acids at the C-terminal of the α-subunit (Belfiore, Malaguarnera et al., 2017, Hernandez-Sanchez, Mansilla et al., 2008). The two isoforms (INSR-A and INSR-B) have distinct biochemical and functional properties and tissue distributions (Belfiore et al., 2017). To first characterize the possible role of INSR signaling in the dystrophic retina, we evaluated the retinal expression of each isoform in both physiological and pathological conditions: retinal *Insr-a* and *Insr-b* expression was analyzed by reverse transcriptase polymerase chain reaction (RT-PCR) in wild type (WT) mice and in the *Pde6b*^*rd10/rd10*^ (*rd10*) mouse model of RP (Chang, Hawes et al., 2007). Retinal RNA extracts were obtained at different stages from postnatal day 15 (P15) (i.e. before the appearance of evident retinal degeneration) up to P60 (after rod loss) (Fig. 1A). The *Insr-a* isoform, which lacks exon 11, was the only isoform detected in both WT and *rd10* retinas at all stages analyzed, as well as in the three adult WT brain areas analyzed (cerebellum, brainstem, remainder of the telencephalon-diencephalon and olfactory bulb). Conversely, key glucoregulatory tissues, such as the liver and adipose tissue, preferentially expressed the *Insr-b* isoform, which contains exon 11.

**Figure 1.**
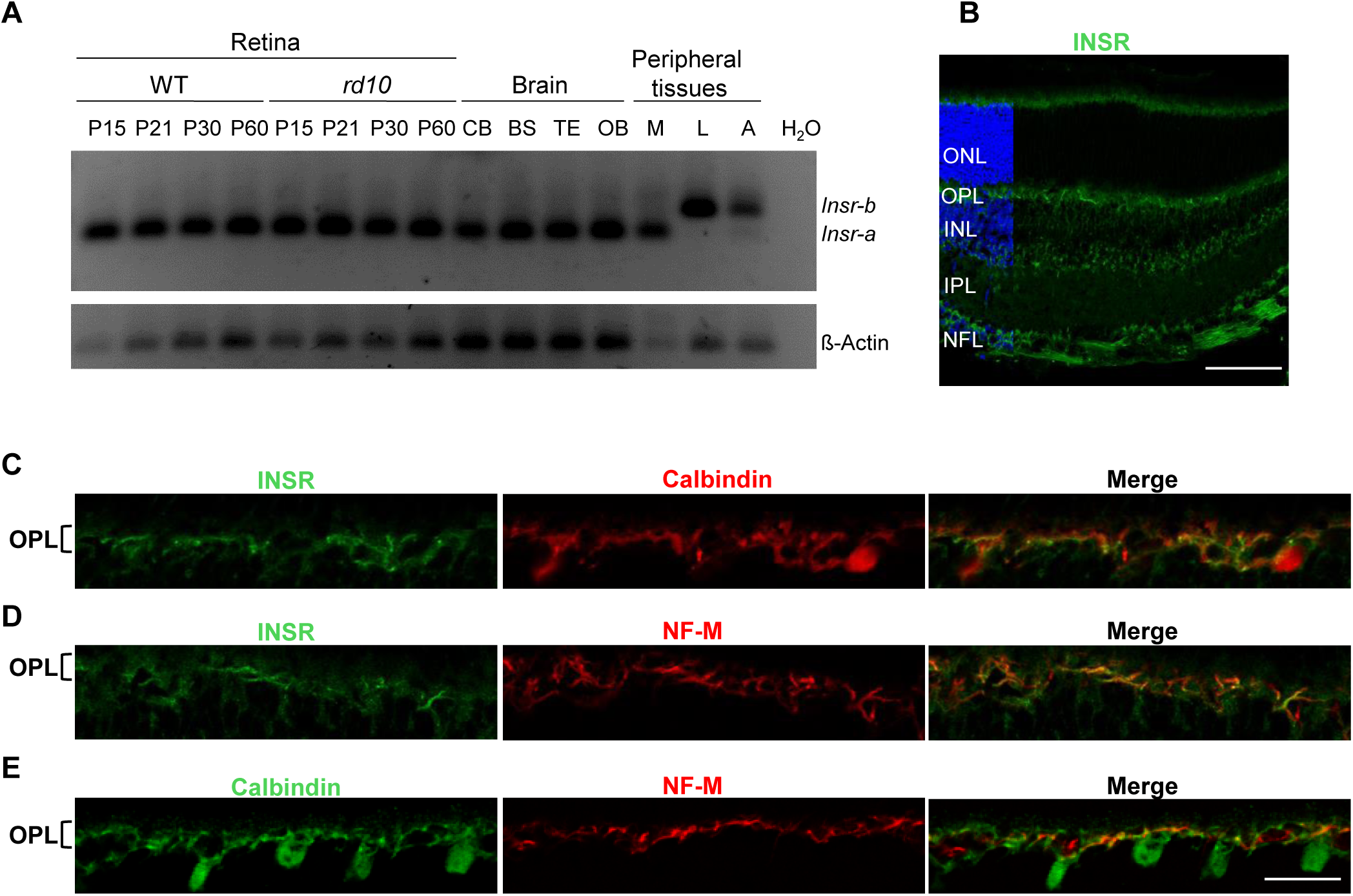
Insulin receptor expression in the retina. **A.** RT-PCR of WT and *rd10* mouse retinas at the indicated ages, and WT adult brain regions and peripheral tissues. Forward and reverse PCR primers corresponded to exons 10 and 12, respectively, of *Insr*. CB, cerebellum; BS, brain stem; TE, remainder of telencephalon and diencephalon; OB, olfactory bulb; M, muscle; L, liver; A, adipose tissue. ß-actin was used as a loading control. **B-E.** Representative images of P21 retinal sections from WT mice. B, Image of a retinal section immunostained for INSR (green). Nuclei are stained with DAPI (blue). **C-E**, Magnified image of the OPL showing double immunostaining for the indicated markers. ONL, outer nuclear layer; OPL, outer plexiform layer; INL, inner nuclear layer; IPL, inner plexiform layer; NFL, nerve fiber layer. Scale bars: 90 μm (B) and 45 μm (C– E).

INSR tissue distribution was visualized by immunofluorescence using the anti-INSR β-subunit antibody C19 (see Table 1). The specificity of this antibody in neural tissue has been previously confirmed in *Insr*-knockout mice (Bruning, Gautam et al., 2000, Dixon-Salazar, Fourgeaud et al., 2014). We found that INSR was widely distributed in the WT retina, in accordance with its versatile role in the CNS. Interestingly, we observed prominent expression in the outer plexiform layer (OPL) and the retinal nerve fiber layer (NFL) (Fig. 1B). The OPL is a synaptic layer in which photoreceptor, horizontal, and bipolar cell axons and dendrites connect. Double immunostaining for INSR and calbindin, a marker of horizontal cells, revealed that INSR expression in the OPL was restricted to a subset of calbindin-positive fibers (Fig. 1C). By contrast, we observed no colocalization of INSR and PKC-α, which labels the dendrites of ON-bipolar cells (Suppl. Fig. 1A). To determine the type (axon or dendrite) of INSR-positive horizontal cell processes, we performed immunostaining for neurofilament M (NF-M), which is selectively expressed by the axons (Peichl & Gonzalez-Soriano, 1993, Peichl & Gonzalez-Soriano, 1994) (Fig. 1E). INSR expression in the OPL was restricted to NF-M-positive horizontal cell axons (Fig. 1D). NF-M-positive ganglion cell axons also showed robust INSR expression (Suppl. Fig. 1B). Horizontal cells receive rod and cone inputs separately; while their axon terminals receive inputs from rods, their dendrites collect input from cone pedicles (Feigenspan & Babai, 2015, Kolb, 1970, Kolb, 1974, Peichl & Gonzalez-Soriano, 1994). Therefore, our results indicate preferential expression of INSR in the horizontal cell axons that synapse with rod spherules.

**Table 1.**

| **Antibody against** | **Host species** | **Dilution** | **Manufacturer** | **Catalog number** |
| --- | --- | --- | --- | --- |
| Calbindin | Mouse | 1:500 | Sigma | C9848 |
| Ctbp2 | Mouse | 1:300 | BD Biosciences | 612044 |
| GluA2 | Mouse | 1:300 | Neuromab | 75-002 |
| Insulin receptor<br>(INSR) | Rabbit | 1:200 | Santa Cruz | C19; SC-711 |
| mGluR6 | Guinea Pig | 1:200 | Acris GmBh | AP20134SU-N |
| Neurofilament<br>Medium (NF-M) | Mouse | 1:500 | Developmental<br>Studies<br>Hybridoma Bank | DH-3 |
| PKC-α | Mouse | 1:500 | Abcam | AB11723 |
| pS6<sup>(Ser240/Ser244)</sup> | Rabbit | 1:200 | Cell Signaling | 5364 |
| Ribeye | Rabbit | 1:400 | Synaptic Systems | 192103 |

Next, we sought to characterize the pattern of INSR expression associated with retinal dystrophy. In early disease stages, before the onset of any significant disturbances in rod cells (P16), comparable global patterns of INSR immunostaining were observed in WT and *rd10* retinas (Suppl. Fig. 2A). However, during the course of rod degeneration (P21–P23) we observed a selective and progressive decrease in INSR immunostaining in horizontal cell axons in the *rd10* retina (Fig. 2). This decrease was not due to the loss of horizontal cells, since similar numbers of calbindin-positive cells were observed in WT and *rd10* retinas at these ages (Suppl. Fig. 3), nor due to degeneration of horizontal cell axons, as evidenced by the preservation of NF-M immunostaining (Fig. 2). Moreover, INSR downregulation was specific to horizontal cell axons; we observed no significant changes in INSR immunostaining in ganglion cell axons (Fig. 2).

**Figure 2.**
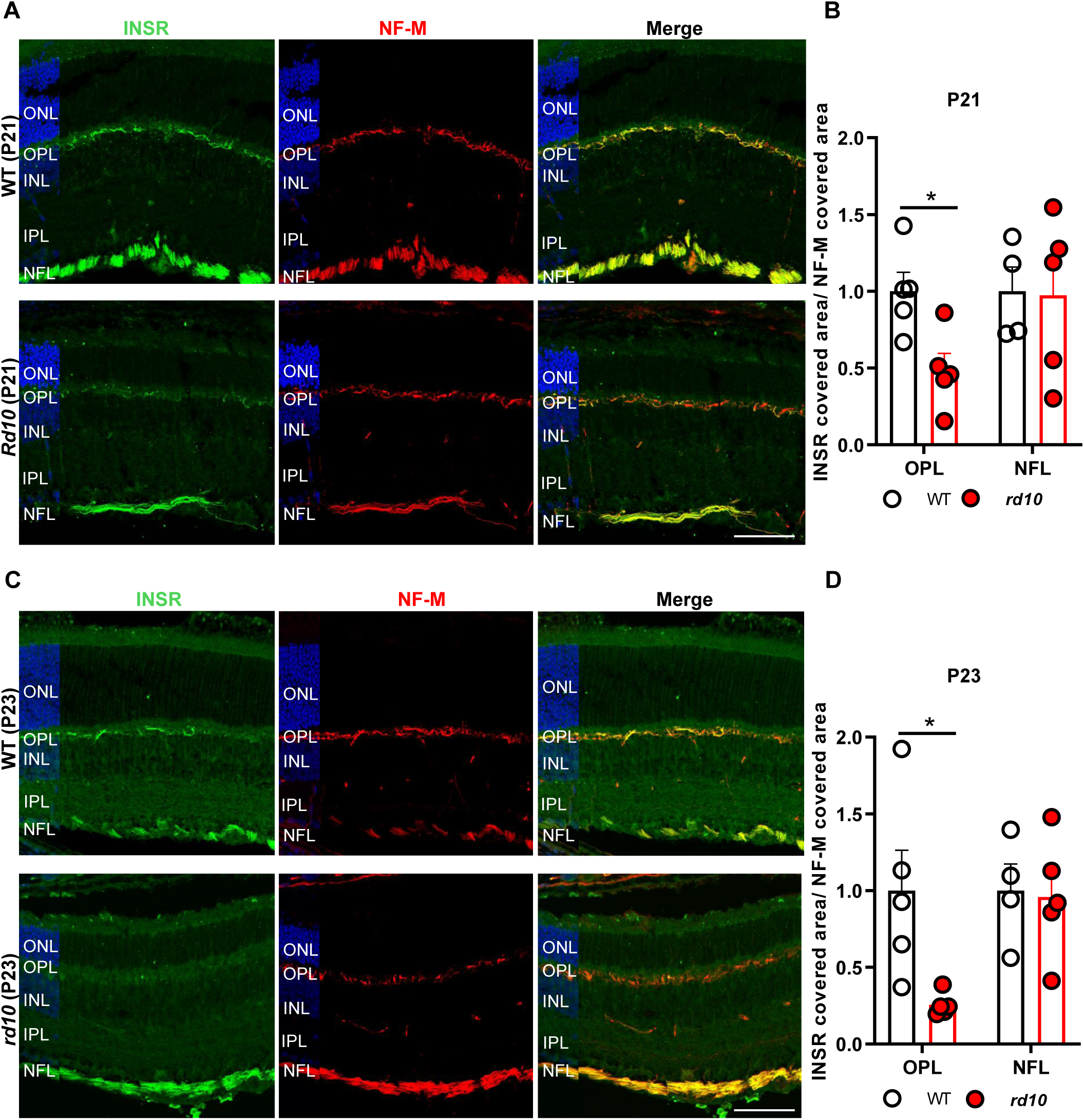
Downregulation of insulin receptor expression in the *rd10* retina. **A, C.** Representative images of P21 (A) and P23 (C) retinal sections from WT and *rd10* mice co-immunostained for INSR (green) and neurofilament-M (NF-M, red). Nuclei are stained with DAPI (blue). ONL, outer nuclear layer; OPL, outer plexiform layer; INL, inner nuclear layer; IPL, inner plexiform layer; NFL, nerve fiber layer. Scale bar: 66 μm. **B, D.** Quantification of the area of INSR-positive staining in P21 (B) and P23 (D) WT and *rd10* retinal sections. The area corresponding to INSR immunostaining was normalized to that of neurofilament-M within the same region to correct for potential variations among retinal sections, and to *INSR* immunostaining in corresponding WT sections (=1.0). Data are presented as the mean + SEM. n=4-5 mice, 4 images per retina. \**p* ≤ 0.05 (unpaired-T test in B and unpaired-T test with Welch’s correction in D).

We next investigated whether the selective decrease in INSR expression in horizontal cell axons had consequences at the level of local INSR signaling. Of the multiple downstream effectors of INSR signaling, we specifically focused on ribosomal protein S6. A recent study (Agostinone, Alarcon-Martinez et al., 2018) reported robust pS6^Ser240/244^ immunostaining in retinal horizontal and ganglion cells, where we observed prominent INSR expression. Immunostaining of WT retinas for pS6^Ser240/244^ corroborated the aforementioned labeling of horizontal and ganglion cell bodies (Fig. 3A, upper panels and Suppl. Fig. 4). Moreover, a closer examination of pS6^Ser240/244^ staining in the OPL revealed profuse punctate labeling in close apposition to the horizontal cell terminal tips (Fig. 3). Interestingly, pS6^Ser240/244^ immunostaining in the *rd10* retina revealed similar staining of horizontal cell bodies, but a dramatic decrease in punctate labeling (Fig. 3A). Double immunostaining for pS6^Ser240/244^ and calbindin showed a decrease in the number of calbindin-positive tips due to retraction of horizontal terminal fibers caused by photoreceptor loss (Cuenca et al., 2014, Pfeiffer et al., 2020). Moreover, in the *rd10* retina half of the remaining horizontal cell terminal tips were devoid of pS6^Ser240/244^ labeling (Fig. 3A, lower panels and Fig. 3B).

**Figure 3.**
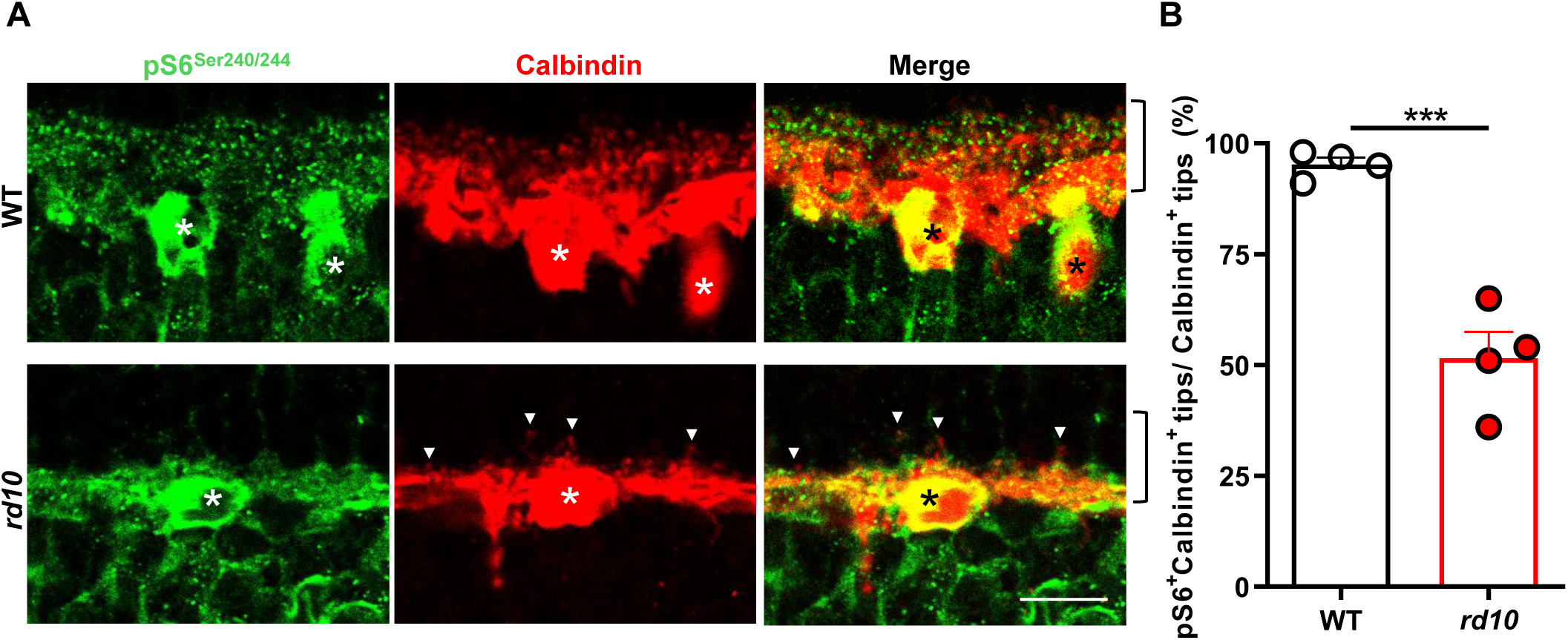
Downregulation of insulin receptor signaling in the *rd10* retina. **A.** Representative images of the OPL of P23 retinal sections from WT and *rd10* mice co-immunostained for pS6^Ser240/244^ (green) and calbindin (red). Arrows indicate horizontal cell terminal tips (calbindin^+^ puncta) without apposite pS6^Ser240/244^ labeling. Asterisks indicate horizontal cell bodies. Brackets delimit the OPL. Scale bar: 12 μm. **B.** Quantification of the number of horizontal terminal tips (calbindin^+^ puncta) with apposite pS6^Ser240/244^ labeling. Data are presented as the mean + SEM. n=4 mice, 4 images. Over 200 calbindin^+^ tips were scored per retina. ***p≤0.001 versus WT (unpaired T-test).

Taken together, our results indicate that in parallel with the degeneration of rod photoreceptors, the terminals of their post-synaptic horizontal cell partners undergo a putative process of insulin resistance, as evidenced by the decreases in INSR levels and pS6^Ser240/244^ signaling. This process may resemble the central and peripheral insulin resistance that occurs in brain neurodegenerative conditions and in type 2 diabetes (Arnold et al., 2018, Holscher, 2020).

### Analysis of synaptic ultrastructure in the OPL of WT and rd10 retinas

Given the marked downregulation of INSR expression and signaling in horizontal cell axons and terminal tips that accompanies rod degeneration, together with the role of INSR in synapse formation and maintenance (Chiu & Cline, 2010, Gralle, 2017, Lee et al., 2011), we next investigated whether changes in INSR expression coincided with alterations in synaptic structure in the OPL. Our analyses focused on rod synapses, since cone survival remains uncompromised in the early stages of degeneration in the *rd10* retina (Zhao, Zabel et al., 2015). The OPL contains so-called triad synapses, formed by the presynaptic rod spherule, two lateral horizontal, and one or two central bipolar postsynaptic terminals invaginating in close apposition to the synaptic ribbon (Fig. 4A and C). Using electron microscopy, we examined WT and *rd10* retinas in the early stages of degeneration (P21), when a large proportion of rod photoreceptor cells persist but signs of degeneration are also evident (Suppl. Fig. 5). In WT retinas, most of the rod synapses (70%) corresponded to characteristic triad synapses (Fig. 4A and D). Conversely, in *rd10* retinas at the same timepoint, triad synapses accounted for less than 20% of all synapses (Fig. 4D). Moreover, in *rd10* retinas more than 40% of rod terminals lacked any postsynaptic element while in WT retinas this was observed in only 8% of rod terminals (Fig. 4B and D). Despite the absence of postsynaptic partners, disconnected rod spherules had a viable appearance, as evidenced by intact mitochondria and electro-lucent content. In these disconnected spherules the synaptic ribbon, when present, was attached to an unusually flat cell membrane with no signs of any postsynaptic invagination (Fig. 4B). By contrast, the degenerative rod spherules (accounting for 20% and 3% of rod terminals in *rd10* and WT retinas, respectively) were highly vacuolated, with electro-dense content and aberrant mitochondria (Fig. 4B and D). Among the degenerated rod spherules we observed both disconnected terminals and triad synapses (Fig. 4B), suggesting that disconnection is not a necessary step before degeneration. Furthermore, both genotypes showed similar numbers of dyads (Suppl. Fig. 6), most likely due to the misalignment of one of the postsynaptic elements with the plane of section (Wang, Pahlberg et al., 2019).

**Figure 4.**
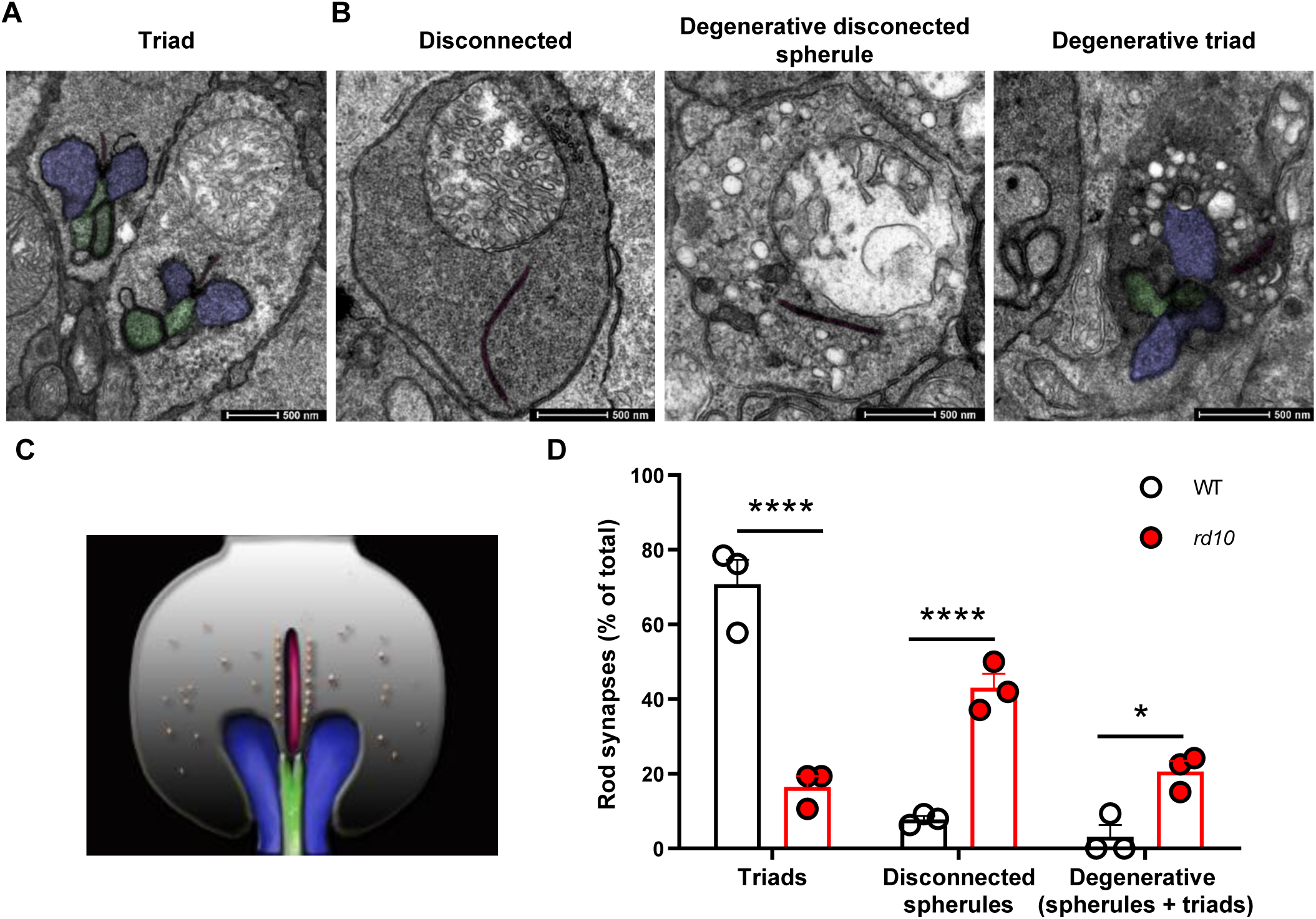
Ultrastructural analysis of rod photoreceptor synapses. **A, B.** Representative electron microscopy images of P21 WT (A) and *rd10* (B) rod synapses. Synapses were categorized as follows: *Triad*, rod spherule with horizontal and bipolar invaginating postsynaptic terminals; *Disconnected*, viable rod spherule without postsynaptic elements; *Degenerative disconnected spherule*, rod spherule lacking postsynaptic profiles with characteristic signs of degeneration; *Degenerative triad*, rod spherule containing postsynaptic profiles but showing characteristic signs of degeneration. Scale bar: 500 nm. **C.** Schematic diagram showing the structure of a rod triad synapse. The rod spherule (grey) consists of the synaptic ribbon (dark red) and a scaffold structure containing presynaptic vesicles (orange circles). Horizontal cell postsynaptic terminals (blue) and bipolar dendrites (green) invaginate into the rod presynaptic terminal (spherule). **D.** Quantification of the proportions of the different synaptic types. Results are expressed as the mean + SEM. n=3 mice. Over 60 synapses were scored per mouse. ****p ≤0.0001, *p≤0.05 (2-way ANOVA followed by Sidak’s multiple comparison test).

Immunostaining using specific markers for photoreceptor ribbon (Ribeye and Ctbp2) and horizontal (GluA2) and bipolar (mGluR6) postsynaptic terminal tips also revealed synaptic disconnection of rod photoreceptors (Suppl. Fig. 7).

### Systemic proinsulin reaches the retina and exerts no metabolic effect

The results described above suggest a possible link between deficient INSR signaling, synaptic disconnection in rod photoreceptors, and the visual impairment characteristic of RP. To determine whether deficient INSR signaling is a disease-modifying factor in RP, and therefore a potential therapeutic target, we employed a gene therapy strategy to enhance INSR signaling. Proinsulin, the insulin precursor, is an INSR-A selective ligand (Malaguarnera, Sacco et al., 2012) with a low metabolic profile, likely due to its poor affinity for the INSR-B isoform (Belfiore et al., 2017, Malaguarnera et al., 2012). We have previously demonstrated the neuroprotective potential of Pi during retinal development and degeneration (Corrochano et al., 2008, Fernandez-Sanchez et al., 2012, Hernandez-Sanchez, Mansilla et al., 2006, Isiegas et al., 2016). Given that treatment of RP in humans would entail long-term administration, we selected Pi to activate INSR-A, the predominant INSR isoform expressed in the retina (Fig. 1A).

To achieve sustained production of Pi, we built upon our previous experience with a recombinant AAV2/1 expressing the human proinsulin (hPi) coding region (Fig. 5; AAV-hPi) (Corpas, Hernandez-Pinto et al., 2017, Fernandez-Sanchez et al., 2012). We first evaluated hPi production following intramuscular administration of AAV-hPi. *rd10* mice received a single injection of AAV-hPi into the gastrocnemius muscle at P10 to induce hPi production before the onset of degeneration (around P18). In AAV-hPi-treated *rd10* mice hPi serum levels were uniformly sustained within each individual mouse for up to 5 months (the maximum follow-up period) (Fig. 5B). Moreover, hPi was detected in whole eye and in retinal extracts as early as 1 week post-injection and up to 7 weeks post-injection (the maximum follow-up period) (Suppl. Fig. 8A). Conversely, hPi was not detected at any timepoint in serum samples, whole eye, or retinal extracts from mice injected with the control vector (AAV-null). Moreover, mature human insulin was absent in serum samples from AAV-hPi-treated WT mice, in agreement with our previous findings (Corpas et al., 2017, Fernandez-Sanchez et al., 2012). Therefore, Pi produced from AAV-hPi remained mainly, if not completely, unprocessed. Importantly, we observed no differences in glycaemia or body weight in AAV-hPi-versus AAV-null-treated mice (Fig. 5C and D), confirming an absence of significant metabolic effects of systemic hPi production, in line with previous studies (Corrochano et al., 2008; Fernández-Sánchez et al., 2012; Corpas et al. 2017).

**Figure 5.**
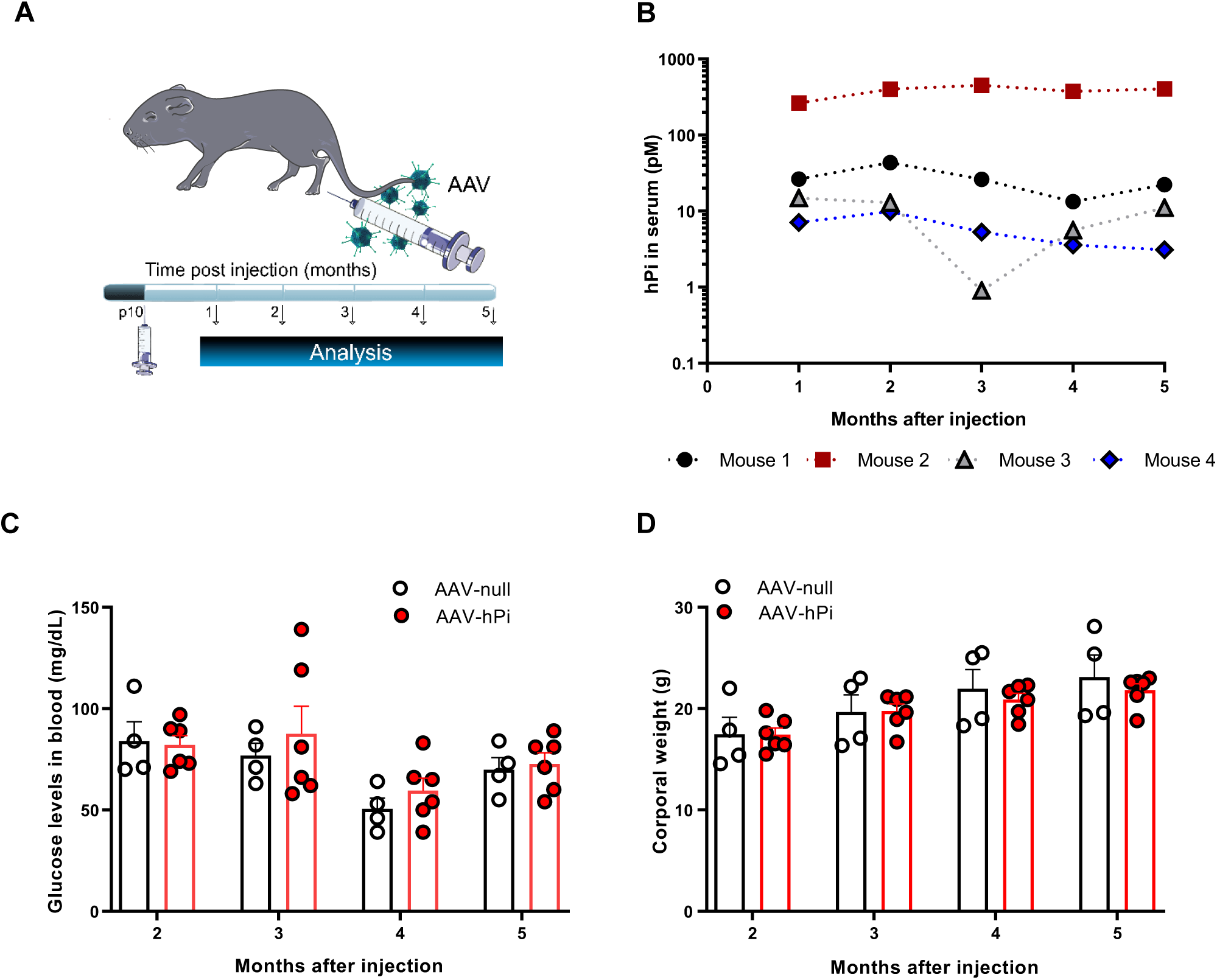
AAV-hPi treatment results in sustained hPi production. **A.** *rd10* mice received a single intramuscular injection of AAV-hPi or AAV-null at P10 and were analyzed at different timepoints post-injection. **B.** Long-term monitoring of serum hPi levels as determined by ELISA in individual *rd10* mice injected with AAV-hPi. **C, D**. Long-term monitoring of glucose levels (C) and body weight (D) in AAV-hPi or AAV-null injected *rd10* mice. Data are presented as the mean + SEM. n=4 mice in B and 4–6 mice per group in C and D.

### AAV-hPi treatment in rd10 mice preserves photoreceptors and photoreceptor synaptic connectivity

To determine the effect of hPi on the dystrophic retina, we first examined whether hPi treatment could restore INSR signaling. AAV-hPi treated *rd10* retinas exhibited higher levels of punctate pS6^Ser240/244^ staining in the OPL than control (AAV-null-treated) *rd10* retinas (Fig. 6). Quantification of the number of pS6^Ser240/244^-positive horizontal cell tips at P21 showed that hPi treatment increased the proportion of calbindin-pS6^Ser240/244^-positive tips to near 90%, close to the proportion observed in WT retinas (Fig. 3). By contrast, in control (AAV-null-treated) *rd10* retinas the proportion of pS6^Ser240/244^-positive horizontal cell tips remained in the range of 50–60% (Fig. 6B). In addition, we observed a greater abundance of total calbindin-positive tips in the AAV-hPi-treated than in the control retinas, most likely a consequence of photoreceptor preservation caused by hPi treatment (Fig. 7).

**Figure 6.**
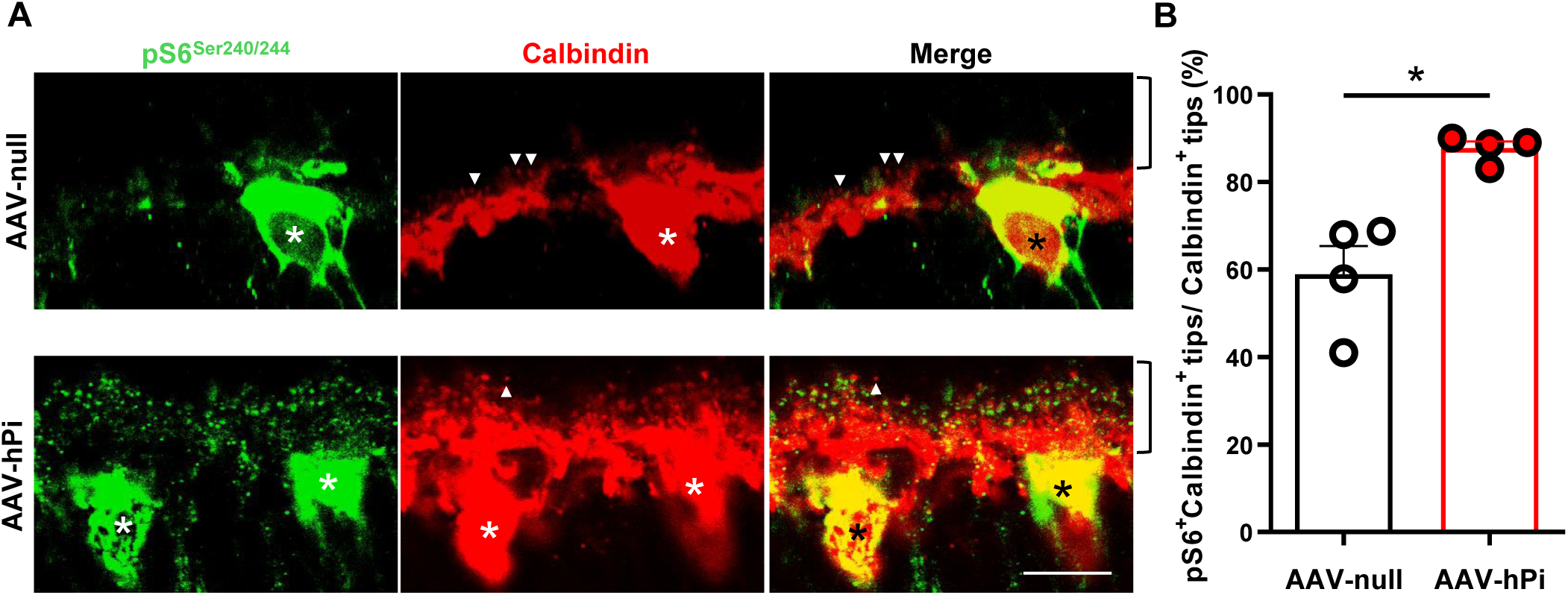
Effect of AAV-hPi administration on insulin receptor signaling. *rd10* mice received a single intramuscular injection of AAV-null or AAV-hPi at P12 and pS6^Ser240/244^ expression was analyzed at P21. **A.** Representative images of the OPL in P21 retinal sections co-immunostained for pS6^Ser240/244^ (green) and calbindin (red). Arrows indicate horizontal cell terminal tips (calbindin^+^ puncta) without apposite pS6^Ser240/244^ labeling. Asterisks indicate horizontal cell bodies. Brackets delimit the OPL. Scale bar: 11 μm. **B.** Quantification of the number of horizontal terminal tips (calbindin^+^ puncta) with apposite pS6^Ser240/244^ labeling. Data are presented as the mean + SEM. n=4 mice, 4 images. Over 200 calbindin^+^ tips per retina were analyzed. *p≤0.05 versus AAV-null (unpaired T-test with Welch’s correction).

**Figure 7.**
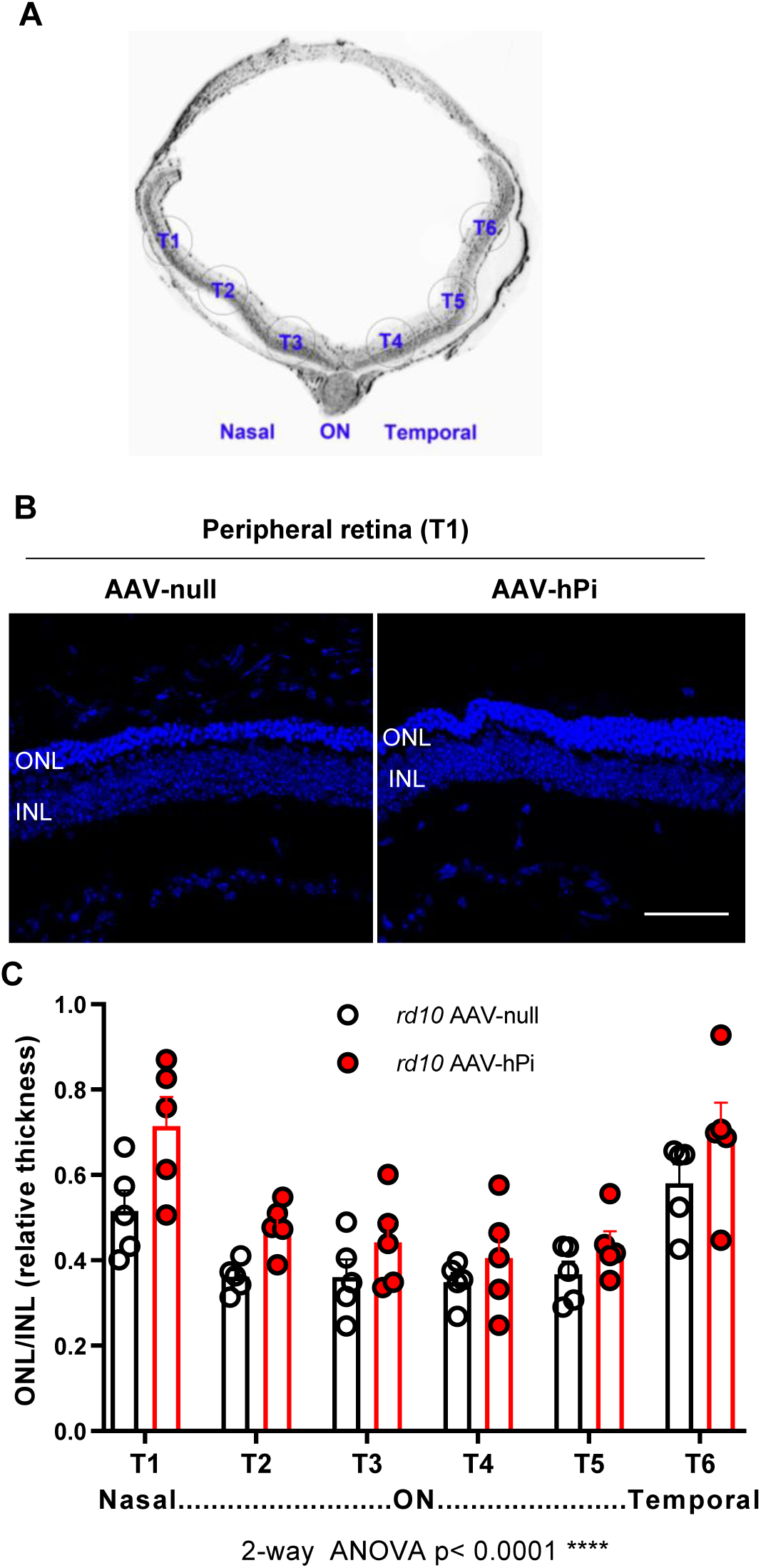
Effect of AAV-hPi administration on photoreceptor preservation. *rd10* mice received a single intramuscular injection of AAV-null or AAV-hPi at P12 and were analyzed at P30. **A.** Schematic depicting retinal sections. The 6 retinal zones in which quantification was performed (T1, T2, T3, T4, T5 and T6) are indicated. ON, optic nerve. **B.** Representative images of P30 retinal sections showing the T1 region from AAV-null- and AAV-hPi-treated *rd10* mice. Blue color corresponds to DAPI staining. ONL, outer nuclear layer; INL, inner nuclear layer. Scale bar: 70 μm. **C.** ONL and INL thickness were measured in equatorial sections corresponding to 6 retinal areas, following a nasotemporal sequence (T1–T6 as defined in panel A). Plot shows the mean + SEM. n=5 mice, 3 sections per retina, 6 regions, 3 measurements per region. ON, optic nerve. ****p ≤0.0001 (2-way ANOVA).

We next investigated whether long-term hPi treatment exerted a neuroprotective effect. To this end, we injected *rd10* mice with AAV-null or AAV-hPi at P10–P12 and analyzed the corresponding retinas at P30, at which point most photoreceptor cells have been lost in this mouse model. Two distinct histological parameters were evaluated: photoreceptor cell preservation and synapse maintenance. AAV-hPi-treated *rd10* retinas showed modest but significant photoreceptor preservation, as determined by measuring the relative increase in thickness of the outer nuclear layer (ONL), an effect that was more evident in the nasal peripheral retina (T1; Fig. 7). In parallel, the synaptic connectivity of photoreceptors was assessed by immunostaining rod-bipolar and rod-horizontal synapses. As expected, photoreceptor preservation in AAV-hPi-treated retinas led to an increase in the number of ribbons. Moreover, hPi reduced the proportion of disconnected postsynaptic terminals of rod cells (i.e. those lacking either a GluA2-positive horizontal postsynaptic terminal [Fig, 8A and C] or a mGluR6-positive bipolar postsynaptic terminal [Fig. 8B and D]). These results reveal a novel effect of proinsulin: preservation of the synaptic connectivity of rod cells with their postsynaptic second-order neuronal partners. This observation suggests a second distinct role of hPi that may be partially independent of its effects on photoreceptor cell survival revealed in our previous studies (Corrochano et al., 2008, Fernandez-Sanchez et al., 2012, Isiegas et al., 2016). Moreover, the increase in the number of photoreceptor synapses induced by proinsulin treatment that we previously observed in the P23H rat model of RP (Fernandez-Sanchez et al., 2012) is most likely a consequence of the rescue of photoreceptor cells as well as preservation of their synaptic contacts, as described here.

**Figure 8.**
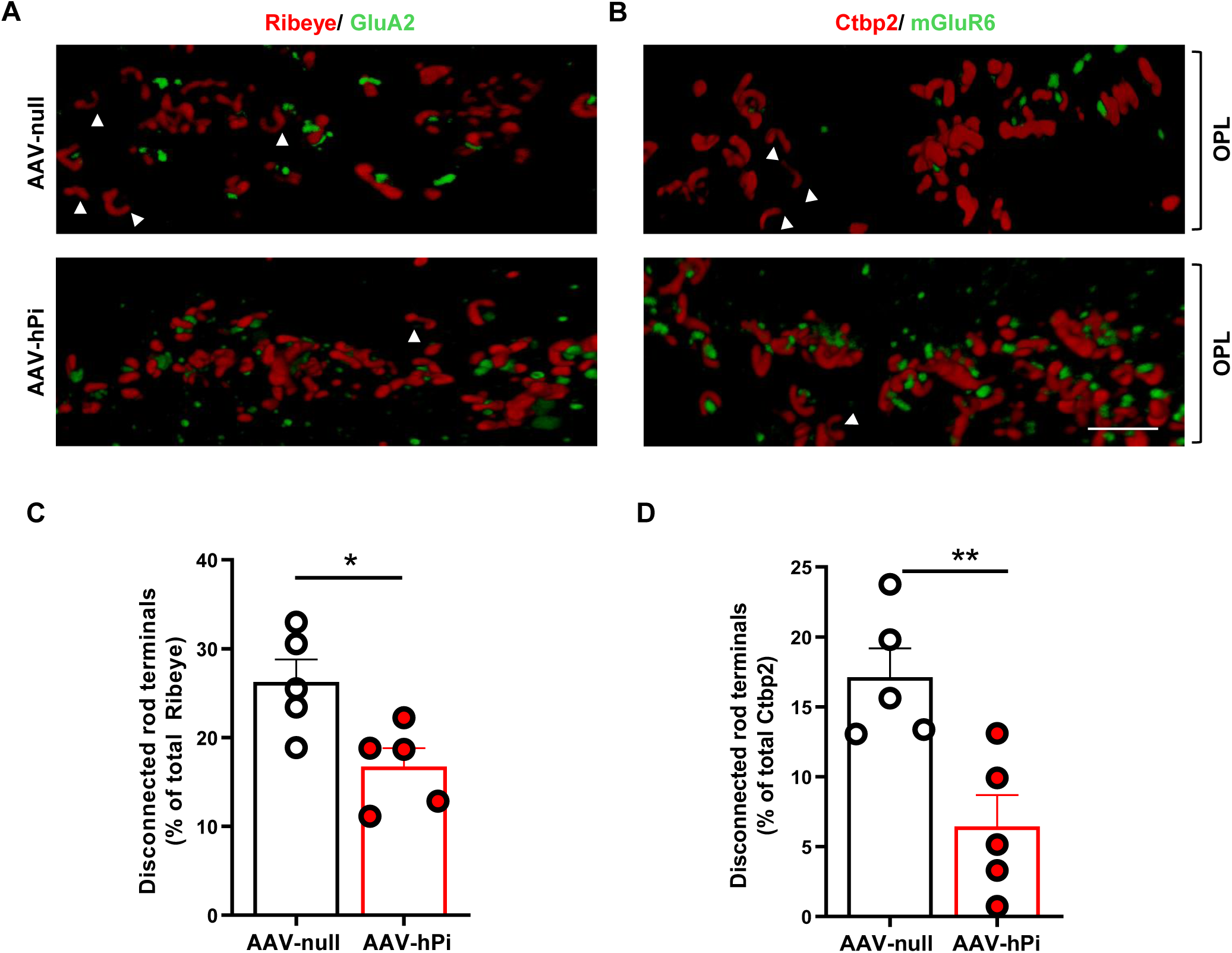
Effect of AAV-hPi administration on rod horizontal and rod bipolar synapses. *rd10* mice received a single intramuscular injection of AAV-null or AAV-hPi at P12 and were analyzed at P30. **A, B.** Representative 3D reconstructions of synapses in the OPL. Retinal sections were co-immunostained for Ribeye (ribbon at presynaptic terminal, red) and the glutamate receptor subunit GluA2 (horizontal postsynaptic terminal, green) (A), or for Ctbp2 (ribbon at presynaptic terminal, red) and the glutamate receptor mGluR6 (bipolar postsynaptic terminal, green) (B). Arrowheads indicate ribbons (rod presynaptic terminals) without post-synaptic partners. OPL, outer plexiform layer. Scale bar: 3 μm. **C, D**. Quantification of the number of disconnected rod presynaptic terminals. Percentage of ribbons without associated GluA2 (C) or mGluR6 (D) punctate staining. Plots show the mean + SEM. *n* = 5 mice. Over 200 ribbons were analyzed per retina. \**p* ≤ 0.05 (unpaired T-test).

### AAV-hPi treatment in rd10 mice preserves visual function

Finally, in AAV-hPi-treated *rd10* mice we assessed whether hPi preserved visual function, which is the most clinically relevant outcome for a potential RP treatment. Electroretinographic (ERG) recordings were performed in dim and daylight conditions every 10 days between P30 and P60 to evaluate rod- and cone-mediated light responses (Fig 9). AAV-hPi-treated *rd10* mice displayed better defined and more prominent ERG waves than their AAV-null-treated counterparts (Fig 9B). Waves corresponding to rods (b-scotopic wave), cones (b-photopic wave), and both photoreceptors (a-mixed and b-mixed waves) were of a significantly greater amplitude in AAV-hPi-treated than AAV-null-treated *rd10* mice (Fig. 9C). ERG recordings thus confirmed that hPi treatment preserved visual function, consistent with the aforementioned preservation of photoreceptor cells and their synapses. Optokinetic testing further confirmed partial preservation of the light response in AAV-hPi-treated *rd10* retinas. In addition to measuring the retinal response to light, this visual behavior test evaluates the function of other components of the visual system, namely optic nerve transmission and visual integration in the brain (Abdeljalil, Hamid et al., 2005). Mice instinctively respond to rotating vertical bars with characteristic movement of their heads in the same direction as the rotation of the bar (Fig. 10A and B). AAV-hPi-treated *rd10* mice showed greater contrast sensitivity than AAV-null-treated counterparts at the two ages studied (P40 and P50; Fig. 10C and D). Together, our results support the disease-modifying potential of Pi, which can prolong visual function in an animal model of RP and therefore constitutes a viable candidate therapy for retinal dystrophies.

**Figure 9.**
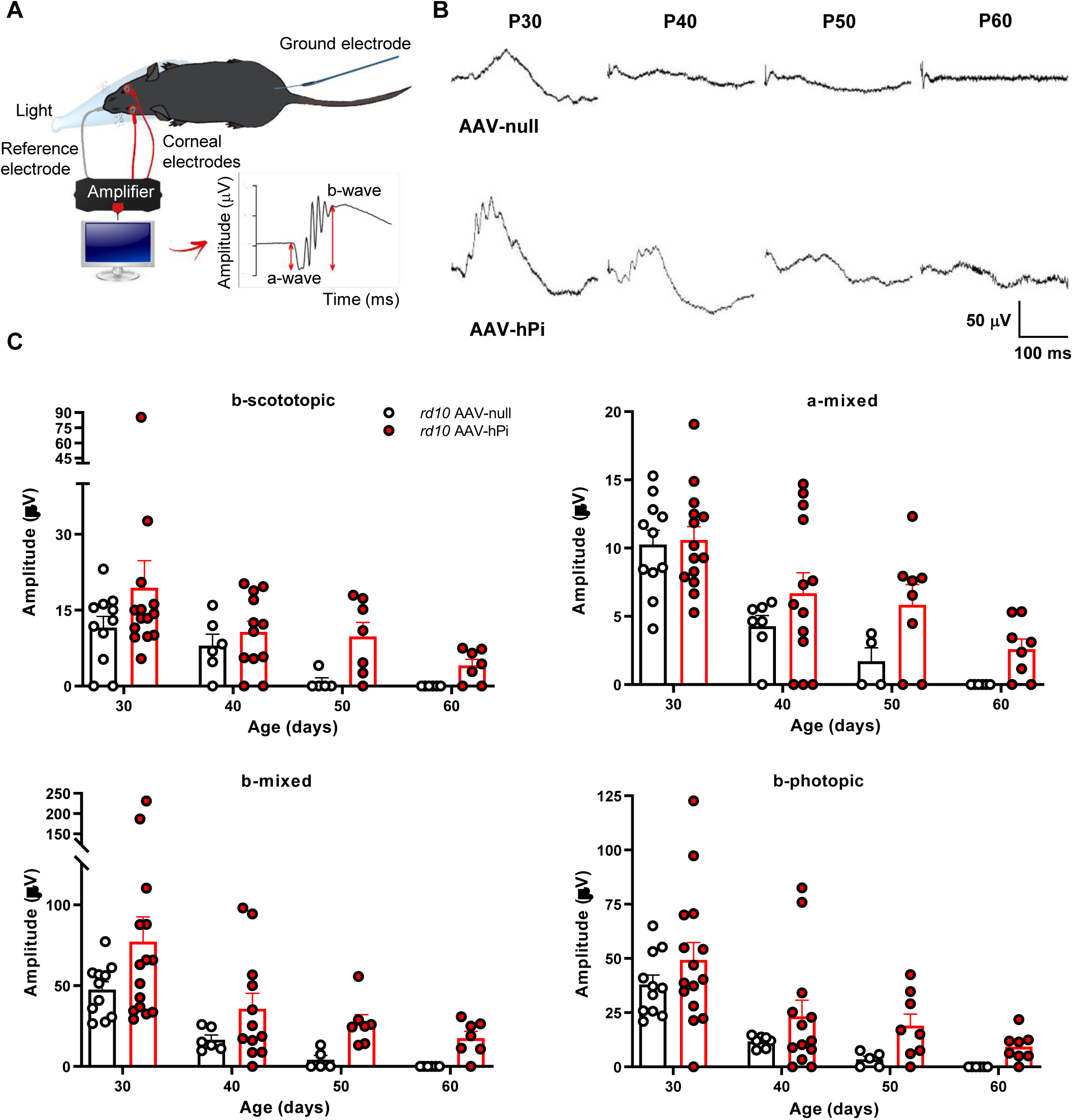
Effect of AAV-hPi administration on retinal response to light. *rd10* mice received a single intramuscular injection of AAV-null or AAV-hPi at P10. ERG recordings were performed at the indicated ages. **A.** Schematic diagram depicting the ERG recording method. **B.** Standard ERG representative trace recordings of the mixed response obtained from 1 AAV-null- and 1 AAV-hPi-treated mouse over the course of the study (P30–P60) in response to a light stimulus of 1.5 cd·s/m^2^. Note the differences in the traced amplitudes between the 2 animals. **C.** Graphs show averaged ERG wave amplitudes, plotted as a function of animal age. Amplitudes of the rod response (b-scotopic; light intensity = −2 log cd·s/m^2^) and rod and cone mixed response (a-mixed and b-mixed; light intensity = 2 log cd·s/m^2^) were recorded under scotopic conditions after overnight adaptation to darkness. Cone amplitudes (b-photopic; light intensity = 2 log cd·s/m^2^) were recorded after 5 minutes of light-adaptation (30 cd/m^2^ background light) under photopic conditions. Results are expressed as the mean + SEM. n=4–13. Significant differences between AAV-null- and AAV-hPi-treated mice were observed for b-scotopic, a-mixed, b-mixed, and b-photopic responses (*p* ≤ 0.05) (2-way ANOVA).

**Figure 10.**
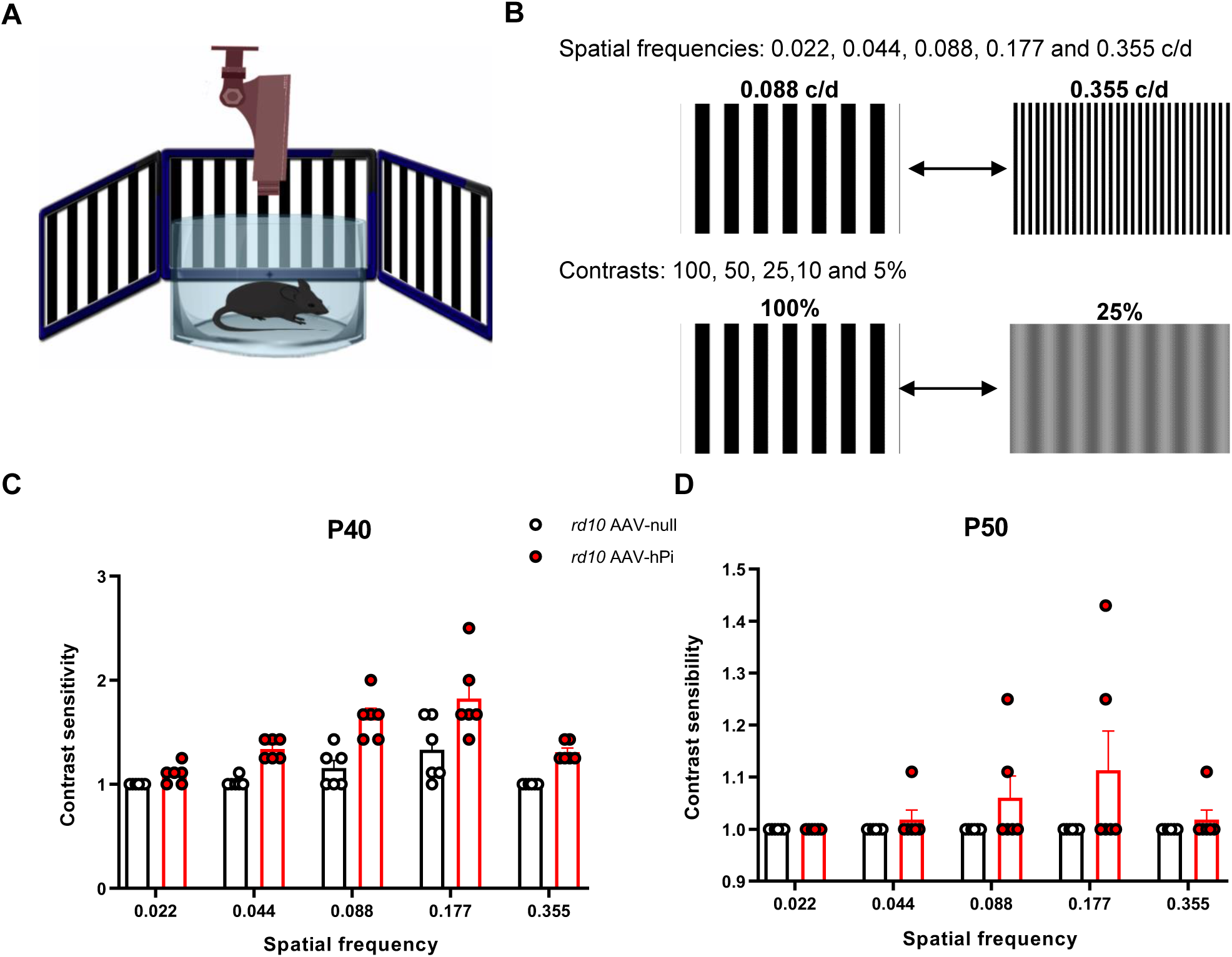
Effect of AAV-hPi administration on optokinetic response. *rd10* mice received a single intramuscular injection of AAV-null or AAV-hPi at P10. The optomotor test was performed at the indicated ages. **A, B.** Schematic depicting the optomotor test (A); 5 different spatial frequencies were tested (0.022–0.355 cycles/degree). The contrast of the moving bars was adjusted (from 100% to 5%) to determine contrast sensitivity (B). **C, D.** Optokinetic responses recorded at the indicated ages in AAV-null- and AAV-hPi treated *rd10* mice. Contrast sensitivity is represented as a function of spatial frequency. Data are presented as the mean + SEM. n=6. Significant differences were observed between AAV-null- and AAV-hPi treated mice at P40 (*p* ≤ 0.05) and P50 (*p*≤0.0001) (2-way ANOVA).

## DISCUSSION

This study describes the downregulation of retinal INSR levels and local signaling during the early stages of retinal neurodegeneration in the *rd10* mouse model of RP, together with concomitant disruption of photoreceptor triad synapses. These findings suggest that deterioration of INSR signaling contributes to the structural and functional degeneration of the retina in *rd10* mice. We provide proof of concept of the neuroprotective effect of INSR stimulation with the insulin precursor Pi, which is selective for the INSR-A isoform found in the retina. Gene therapy using an AAV induced sustained production of circulating Pi, which reached the retina, restored local INSR signaling, and exerted neuroprotective effects on retinal structure and visual function, without affecting peripheral metabolic parameters. Proinsulin treatment attenuated both photoreceptor cell loss and synaptic disconnection, and prolonged visual function, highlighting the potential of Pi as a candidate therapy for RP.

Studies over the past two decades have broadened the scope of INSR activity far beyond the peripheral metabolic role initially ascribed to this receptor. The versatility of this receptor is also implied by its widespread expression in the CNS. At the neuronal level, INSR has been implicated in synaptic plasticity, dendritic outgrowth, and cell survival (Chiu & Cline, 2010, Gralle, 2017, Lee et al., 2011). The *Insr-a* splice variant, which is predominantly expressed in different areas of the brain as well as the retina, is the most ancient and promiscuous isoform, and participates in insulin, Pi, and IGF-II signaling (Hernandez-Sanchez et al., 2008, Malaguarnera et al., 2012). Interestingly, many of the signaling pathways in which INSR-A is involved are similar to those mediated by growth factor receptors, and in some ways this isoform resembles the ancestral INSR expressed in invertebrates and low vertebrates (Banks et al., 2012, Belfiore, Frasca et al., 2009, Chan & Steiner, 2000). The present findings confirm the previously described wide distribution of INSR in the WT retina (Gosbell, Favilla et al., 2000, Rajala, Wiskur et al., 2009). However, we also describe for the first time more intense INSR immunostaining in horizontal and ganglion cell axons than in other retinal structures. Interestingly, analysis of INSR expression during retinal degeneration in the rd10 mouse revealed local downregulation of INSR, specifically in horizontal cell axons. This was accompanied by a decrease in the phosphorylation of ribosomal protein S6 in horizontal cell terminals, indicating impaired INSR signaling. Notably, horizontal cell axon terminals receive input from the rod photoreceptors, the main targets of degeneration in RP. A recent study by Agostinone et al. (Agostinone et al., 2018) reported specific decreases in pS6^Ser240/244^ in ganglion cells following transection of their axons, but no alterations in pS6^Ser240/244^ in horizontal cells. In the present study, we observed selective downregulation of INSR expression and impaired signaling in the horizontal cells with which the damaged rod photoreceptors synapse. It remains unclear whether this finding is mechanistically linked to the decreases in pS6^Ser240/244^ described by Agostinone et al (Agostinone et al., 2018). However, our results are consistent with the proposals by other authors that INSR signaling may vary in a neuronal activity-dependent manner (Chiu & Cline, 2010, Clarke, Mudd et al., 1986, Hori, Yasuda et al., 2005), and suggest that INSR expression and signaling in horizontal cells may depend on rod input.

Interestingly, in *rd10* retinas INSR downregulation and consequent impairment of INSR signaling coincided with the presence of disconnected but otherwise apparently viable rod terminals. Alterations in retinal synaptic circuitry as part of retinal remodeling during photoreceptor degeneration have been acknowledged for more than two decades (Jones, Kondo et al., 2012, Lewis, Linberg et al., 1998, Pfeiffer et al., 2020, Strettoi, Pignatelli et al., 2003). De-afferentiation of second-order neurons has typically been described in advanced stages of degeneration as a consequence of photoreceptor loss (Jones et al., 2012, Lewis et al., 1998, Pfeiffer et al., 2020, Strettoi et al., 2003). However, here we describe early disconnection events affecting up to 50% of apparently viable rod spherules during the initial stages of degeneration. Although further studies will be required to clarify the consequences of this disconnection, it does not appear to necessarily precede rod death, as we detected degenerative triads containing the presynaptic rod spherule and horizontal and bipolar terminals. These observations have direct implications for the development of neuroprotective therapies and could open a new line of research into interventions aimed at preserving rod connectivity.

Our findings demonstrate the potential of Pi as a synaptoprotective factor. In *rd10* mice treated with AAV-hPi, systemically produced Pi reached the retina and restored INSR signaling, as determined by measuring pS6^Ser240/244^ levels. Concomitantly, Pi treatment reduced the number of disconnected rod presynaptic terminals, supporting a role of INSR signaling in photoreceptor synaptic connectivity. Our results are in line with those of the seminal study by Chiu et al. 2008 (Chiu & Cline, 2010), who used optic tectal neurons in living *Xenopus* tadpoles to demonstrate that INSR signaling maintains both synaptic contacts and the branches on which they lie. Moreover, in the aforementioned study by Agostinone et al. (Agostinone et al., 2018) stimulation of INSR signaling with insulin promoted regeneration of the dendritic arbors of retinal ganglion cells after axonal injury.

Deficient local INSR signaling associated with RP may be a general feature of neurodegenerative diseases, irrespective of peripheral insulin resistance [reviewed in (Arnold et al., 2018)]. Several studies have described decreased INSR expression and/or attenuation of the activation states of INSR signaling molecules in affected brain regions in patients with Alzheimer’s (Rivera et al., 2005, Steen et al., 2005, Talbot et al., 2012) and Parkinson’s (Moroo, Yamada et al., 1994, Takahashi, Yamada et al., 1996, Timmons, Coakley et al., 2009) diseases. We show that pharmacological stimulation of INSR signaling with Pi has a disease-modifying effect over the course of retinal degeneration, in line with the beneficial effects of INSR stimulation with insulin reported in other neurodegenerative diseases of the brain and retina (Agostinone et al., 2018, Arnold et al., 2018, Holscher, 2020). Moreover, the widespread expression of INSR in the retina suggests that synaptic maintenance promoted by INSR signaling is only one of several retinal processes regulated by INSR. Indeed, we and others have demonstrated the neuroprotective effects of INSR signaling molecules on photoreceptor survival (Punzo, Kornacker et al., 2009, Rajala, Tanito et al., 2008, Sanchez-Cruz, Villarejo-Zori et al., 2018).

Our results provide a novel mechanism to account for our previous descriptions of the neuroprotective effects of Pi on retinal dystrophy (Corrochano et al., 2008, Fernandez-Sanchez et al., 2012, Isiegas et al., 2016) and cognitive impairment (Corpas et al., 2017), and further support the validity of Pi-mediated stimulation of INSR as a candidate therapy for neurodegenerative conditions of the CNS.

## MATERIALS AND METHODS

### Animals

The *rd10* mouse is an autosomal recessive homozygous mutant for phosphodiesterase 6b (*Pde6b*^*rd10/rd10*^) on a C57BL/6J background (Chang et al., 2007). Both *rd10* and WT control mice of the same background were obtained from The Jackson Laboratory (Bar Harbor, ME, USA). All animals were housed and handled in accordance with the ARVO statement for the Use of Animals in Ophthalmic and Vision Research, European Union guidelines, and those of the local ethics committees of the CSIC and the Comunidad de Madrid (Spain). Mice were bred in the CIB core facilities on a 12/12-h light/dark cycle. Light intensity was maintained at 3–5 lx.

### Generation and administration of adeno-associated viral vectors

Recombinant AAV serotype 2/1 viral vectors bearing cDNA from the human proinsulin (hPi) gene under control of the cytomegalovirus promoter (AAV-hPi) or without hPi cDNA (AAV-null) were generated in the Center for Animal Biotechnology and Gene Therapy at the Universitat Autònoma de Barcelona, as previously described (Corpas et al., 2017). *rd10* mice received a single intramuscular injection of 7.2 × 10^11^ vector genomes/kg body weight of AAV-hPi or AAV-null at P10–12. The total vector dose was distributed equally between the gastrocnemius muscles of both hindlimbs.

### Measurement of proinsulin, insulin, and glycaemia

Serum, eye, and retinal levels of human Pi and serum levels of human insulin were measured using human Pi and human insulin ELISA kits (EZHPI-15K and EZHI-14K, respectively; Millipore, Darmstadt, Germany) according to the manufacturer’s instructions and as described previously (Isiegas et al., 2016). Glycaemia was directly measured in blood samples using the Glucocart™ Gmeter kit (A. Menarini Diagnostics Ltd., Berkshire, UK).

### RNA isolation and RT-PCR

Total RNA from tissues was isolated using Trizol reagent (Invitrogen). The reverse transcriptase reaction (RT) was typically performed with 2 µg RNA, the Superscript III Kit, and random primers (all from ThermoFisher Scientific, Waltham, MA), followed by amplification with the 2X PANGEA-Long PCR Master Mix (Canvax Biotech, Córdoba, Spain). Mouse *Insr* was amplified using the following primers: sense, 5’-GGCCAGTGAGTGCTGCTCATGC-3’ (inside exon 10); antisense, 5’-TGTGGTGGCTGTCACATTCC-3’ (inside exon 12). Mouse β-actin was amplified using the following primers: sense, 5’-AAGGCCAACCGTGAAAAGAT-3’; antisense, 5’-GTGGTACGACCAGAGGCATAC-3’.

### Immunofluorescence and image analysis

Animals were euthanized and their eyes enucleated and fixed for 50 minutes in freshly prepared 4% paraformaldehyde in Sörensen’s phosphate buffer (SPB) (0.1 M, pH 7.4), and then cryoprotected by incubation in increasing concentrations of sucrose (final concentration, 50% in SPB). The eyes were then embedded in Tissue-Tek OCT (Sakura Finetec, Torrance, CA, USA) and snap frozen in dry-ice cold isopentane. Equatorial sections (12 µm) were cut on a cryostat and mounted on Superfrost Plus slides (Thermo Scientific, Massachusetts, USA), dried at room temperature, and stored at −20°C until the day of the assay.

Before performing further analyses, slides were dried at room temperature. After rinsing in PBS and permeation with 0.2% (w/v) Triton X-100 in PBS, sections were incubated with blocking buffer (5% normal goat serum, 1% Tween-20, 1 M glycine in PBS) for 1 h and then incubated overnight at 4°C with primary antibodies (Table 1) diluted in blocking buffer. Sections incubated in the absence of primary antibody were used as specificity controls. After rinsing in PBS and incubation with the appropriate secondary antibodies (Table 1), sections were stained with DAPI (4’,6-diamidino-2-phenylindole; Sigma-Aldrich Corp., St. Louis, MO, USA) and cover slipped with Fluoromont-G. For GluA2 and mGluR6 immunostaining, an antigen retrieval step was performed prior to incubation in blocking buffer. To this end, sections were incubated in citrate buffer (10 mM sodium citrate, 0.05% Tween-20, pH 6.0) in boiling water for 10 minutes. After cooling to room temperature, sections were rinsed with PBS and then incubated in freshly prepared 0.2% sodium borohydride in PBS before continuing with the blocking reaction described above.

Sections were analyzed using a laser confocal microscope (TCS SP5 and TCS SP8; Leica Microsystems, Wetzlar, Germany). In all cases, retinal sections to be compared were stained and imaged under identical conditions. To measure the area occupied by INSR and NF-M immunostaining, images were converted to black and white and analyzed with Fiji software. To quantify the number of horizontal cell tips with associated pS6^Ser240/244^ punctate staining, over 200 horizontal cell tips per animal in 4 retinal areas were analyzed. Similarly, for synapse quantification, over 200 rod ribbons per animal in 4 retinal areas were assessed for associated GluA2 (horizontal cell postsynaptic terminal) or mGluR6 (horizontal cell postsynaptic terminal) punctate immunostaining. To evaluate photoreceptor preservation, three sections per retina were analyzed: for each section, six areas in a nasotemporal sequence were photographed (Fig. 7A). ONL and INL thickness were measured in three random positions for each image.

### Transmission electron microscopy

Animals were euthanized and their eyes enucleated and fixed overnight in freshly prepared 2% paraformaldehyde and 2% glutaraldehyde in sodium cacodylate buffer (0.1 M, pH 7.4). After dissection of the cornea and lens, optic cups were post-fixed for 1 hour with 1% OsO_4_ (v/v) and 1% K_3_Fe(CN)_6_ (v/v) in ultrapure water, dehydrated through a graded series of ethanol solutions and embedded in Epoxy EMbed-812 resin (EMS, Electron Microscopy Sciences). Semi-thin (0.5 μm) and ultra-thin sections were obtained using an Ultracut E ultramicrotome (Leica). Semi-thin sections were stained with Toluidine Blue and mounted with Entellan and images were obtained using a light microscope (Axio Observer Z1, Zeiss) with 20× and 100× oil immersion objectives. Ultra-thin sections were contrasted with uranyl acetate and lead citrate, and analyzed at the NUCLEUS electron microscopy facility at the University of Salamanca using a Tecnai Spirit Twin 120 kv electron microscope with a CCD Gatan Orius SC200D camera with DigitalMicrograph™ software. The proportions of the different types of synapses were quantified in ultra-thin sections from 3 mice of each genotype by analyzing over 60 synapses per mouse.

### ERG recordings and optomotor tests

Mice were handled and ERGs performed as previously described (Corrochano et al., 2008). Measurements were performed by an observer blind to experimental condition. ERG signals were amplified and band filtered between 0.3 and 1000 Hz (CP511 Preamplifier, Grass Instruments, Quincy, MA, USA) and digitized to 10 kHz using a PowerLab acquisition data card (AD Instruments Ltd., Oxfordshire, UK). Graphical representations of the signals recorded and luminous stimuli control were performed with Scope v6.4 PowerLab software (AD Instruments, Oxford, UK). ERG wave amplitudes were measured off-line and the results averaged. Wave amplitude analysis was performed using the MATLAB application.

An optomotor device was built based on the design proposed by Prusky et al. (Prusky, Alam et al., 2004). Mice were placed in the center of a square array of computer monitors that displayed stimulus gratings, and then monitored using an overhead infrared television camera placed above the testing chamber. The test started with the most easily visible stimulus, with a spatial frequency of 0.088 cycles/degree, a temporal frequency of 0.88 Hz, and a normalized contrast of 1. Contrast sensitivity was calculated as the inverse of contrast threshold, and was measured at distinct spatial frequencies ranging from 0.022–0.355 cycles/degree (Fig. 10B). The Vision Egg tool was used for light stimulation. Stimuli consisted of vertical black/white bars (gratings) moving through the screens.

### Statistical analysis

Statistical analysis was performed with GraphPad Prism software 8.0 (GraphPad Software Inc., La Jolla, CA, USA).

To compare two groups, normality was assessed for each group using the Shapiro-Wilk normality test. Normally distributed data were analyzed using an unpaired T-test, and non-normally distributed data using a Mann Whitney U-test. Homocedasticity for all data sets was assessed employing F test. When variance between data sets was significantly different Welch’s correction was applied to the T-test. Unpaired T-tests were used to compare insulin receptor and pS6^Ser240/244^ levels between WT and *rd10* animals, and to assess rod-horizontal and rod-bipolar connectivity in AAV-null and AAV-hPi mice.

Analysis of variables over time was achieved using a 1-way ANOVA. Dunnett’s multiple comparison test was used to compare values at different time points with those at a specific time point. Accordingly, serum, eye and retinal hPi levels at different time points were compared with initials values using Dunnett’s multiple comparison test.

Comparisons of two variables were performed using a 2-way ANOVA, followed by Sidak’s multiple comparison test where a significant interaction was detected. Sidak’s multiple comparison test was thus used to compare the different synaptic morphologies between WT and *rd10* animals. Differences in histological parameters (ONL/INL ratio), ERG wave amplitude, and optomotor test results between AAV-null mice and AAV-hPi mice were assessed using a 2-way ANOVA.

In all cases, statistical significance was set at p ≤0.05.

---

**Supplementary Figure 1.**
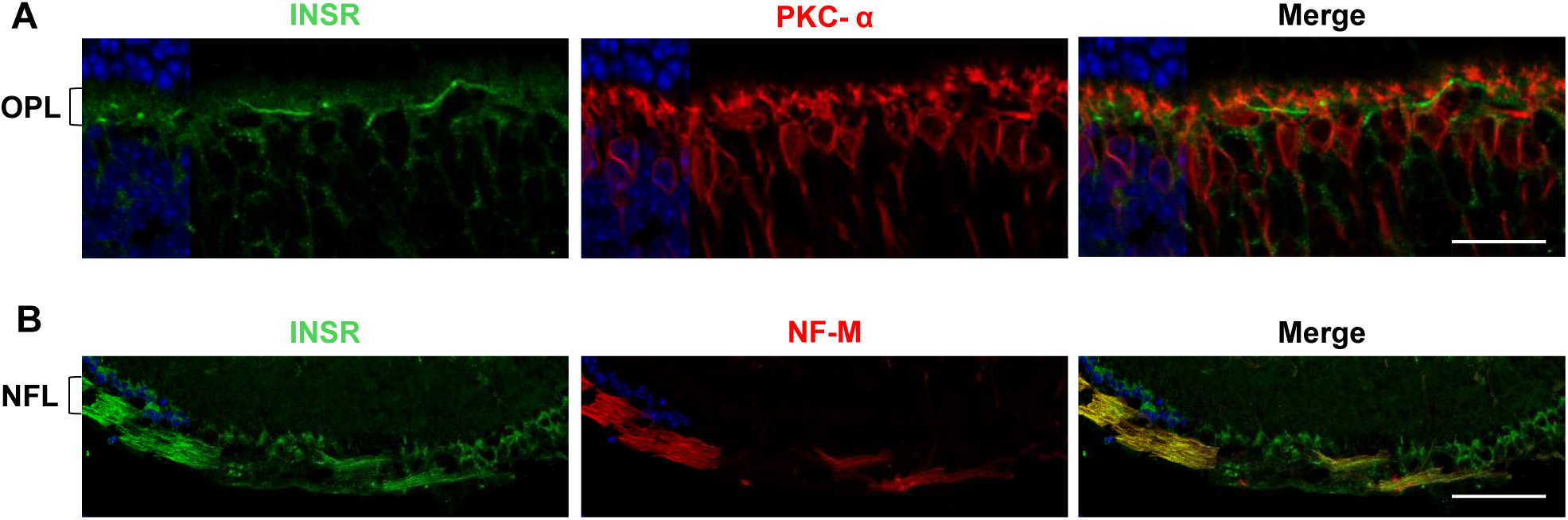
Analysis of insulin receptor expression. Representative retinal sections collected from WT mice at P21. **A, B**. Magnified image showing the OPL (A) and NFL (B) after co-immunostaining for the indicated markers. Nuclei are stained with DAPI (blue). OPL, outer plexiform layer; NFL, nerve fiber layer. Scale bar: 21 μm.

**Supplementary Figure 2.**
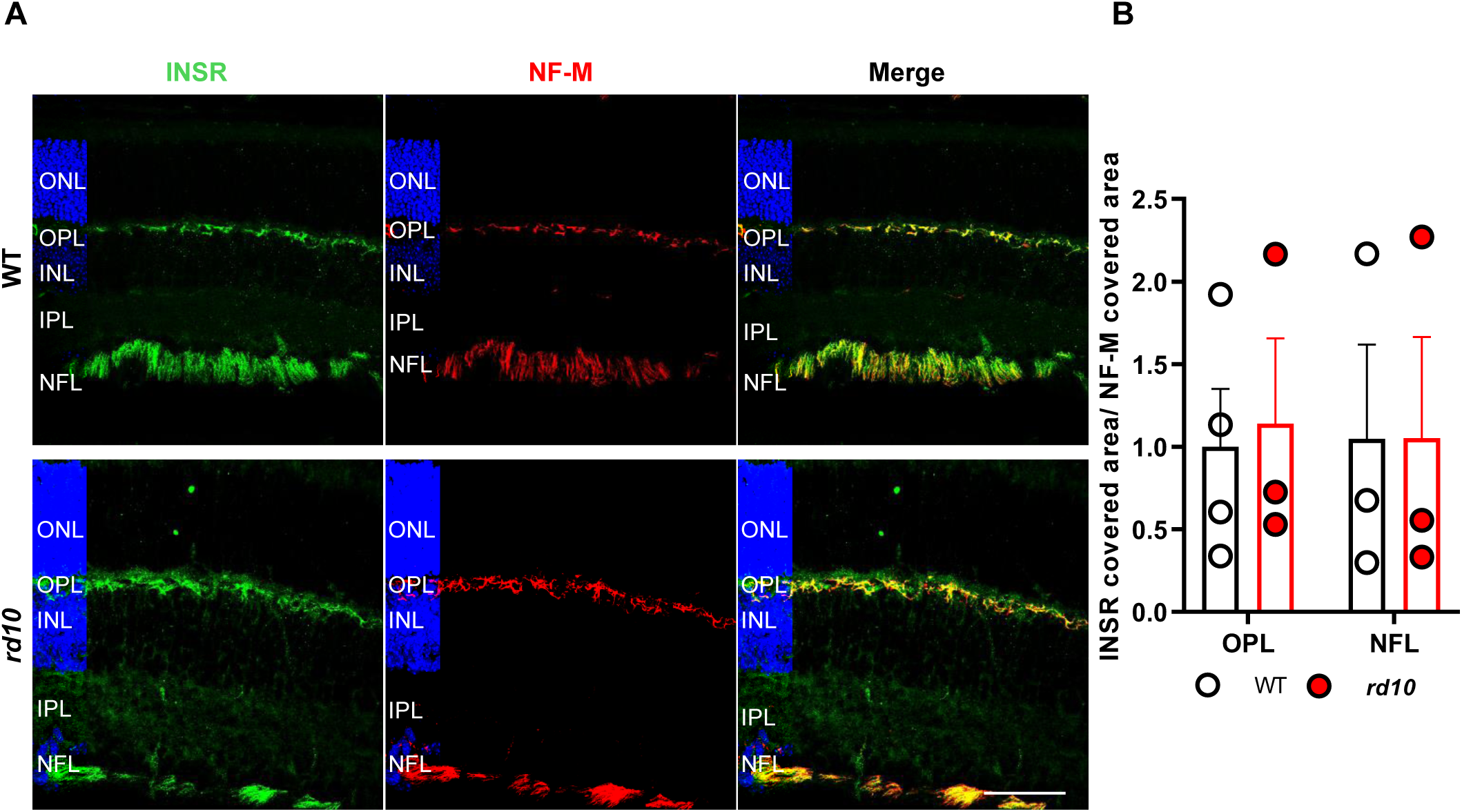
Analysis of insulin receptor expression in WT and *rd10* retinas at P16. **A.** Representative images of P16 retinal sections from WT and *rd10* mice co-immunostained for INSR (green) and neurofilament-M (NF-M, red). Nuclei are stained with DAPI (blue). ONL, outer nuclear layer; OPL, outer plexiform layer; INL, inner nuclear layer; IPL, inner plexiform layer; NFL, nerve fiber layer. Scale bar: 58 μm. **B.** Quantification of the area of INSR-positive immunostaining at P16 in retinal sections from WT and *rd10* mice. The area of INSR-positive immunostaining was normalized to that of neurofilament-staining of the same region to correct for potential variation among retinal sections, and to *INSR* immunostaining in WT sections (=1.0). Data are presented as the mean + SEM. n=3 mice, 4 images per retina.

**Supplementary Figure 3.**
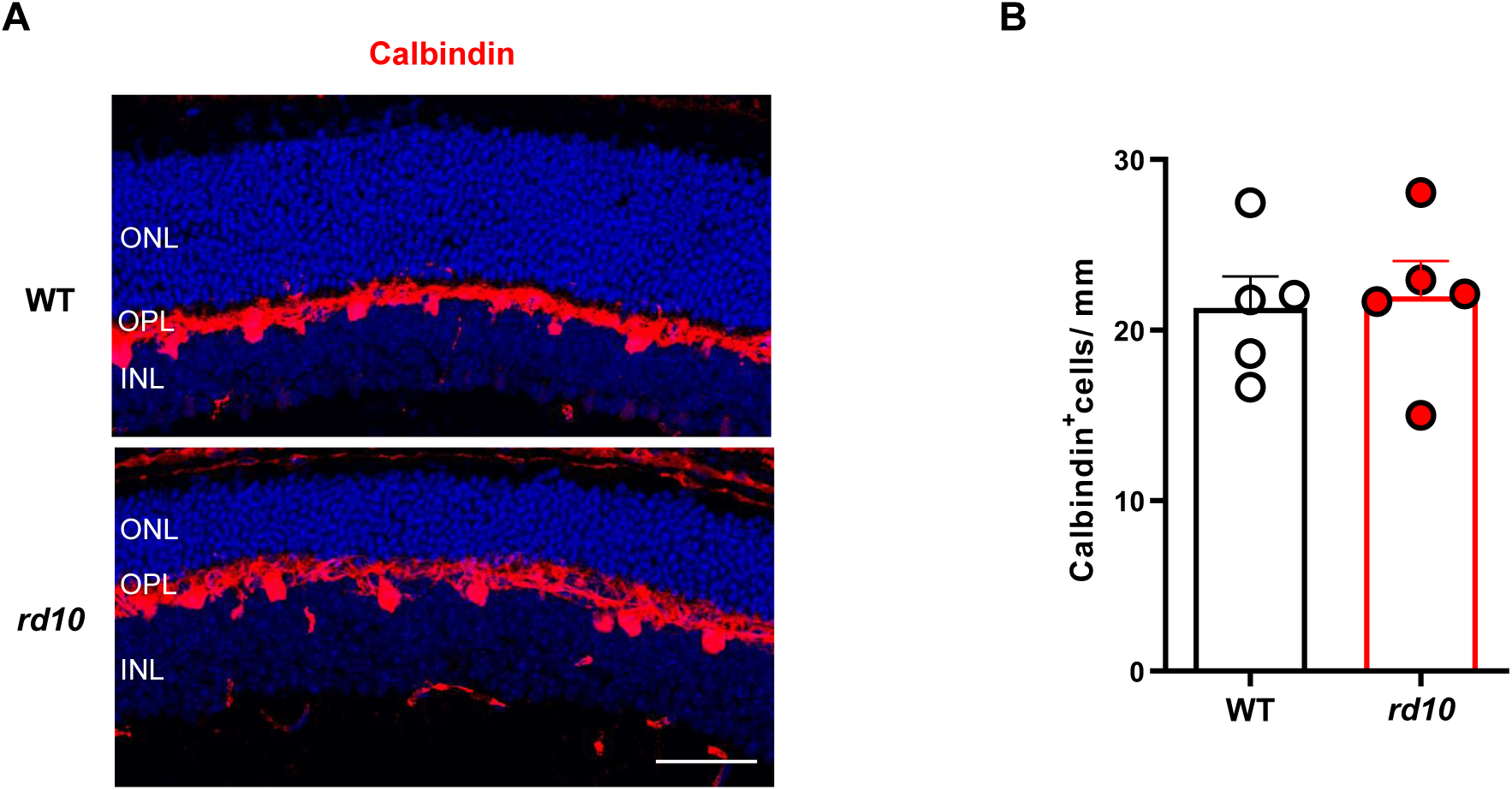
Comparison of the number of horizontal cells in WT and *rd10* retinas. **A.** Representative images of P23 retinal sections from WT and *rd10* mice, immunostained for calbindin to label horizontal cells (red). Nuclei are stained with DAPI (blue). **B.** The number of calbindin^+^ cells was scored in equatorial sections in the central regions of the retina (T3–T4; see Methods and Figure 7A). Plots show the mean + SEM. n=5 mice, 10 images per retina. ONL, outer nuclear layer; OPL, outer plexiform layer; INL, inner nuclear layer. Scale bar: 38 μm.

**Supplementary Figure 4.**
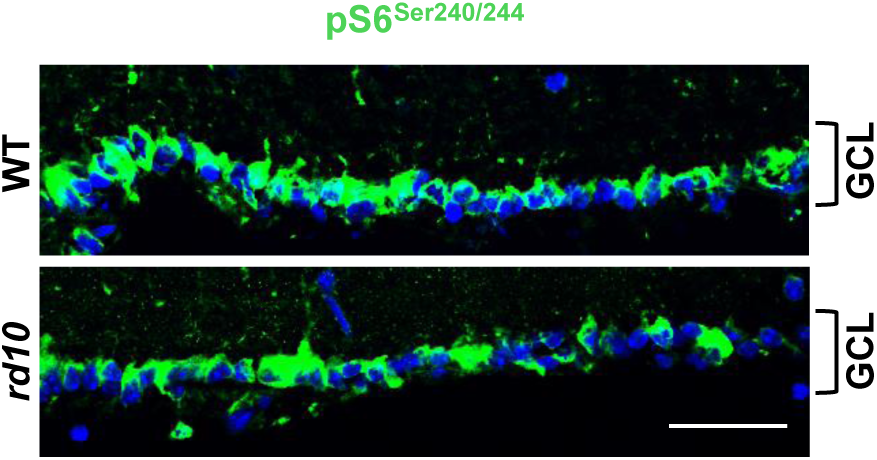
Analysis of pS6^Ser240/244^ expression in ganglion cells in WT and *rd10* retinas. Representative images of the GCL of P23 retinal sections from WT and *rd10* mice immunostained for pS6^Ser240/244^ (green). Nuclei are stained with DAPI (blue). Brackets delimit the GCL. GCL, ganglion cell layer. Scale bar: 41 μm.

**Supplementary Figure 5.**
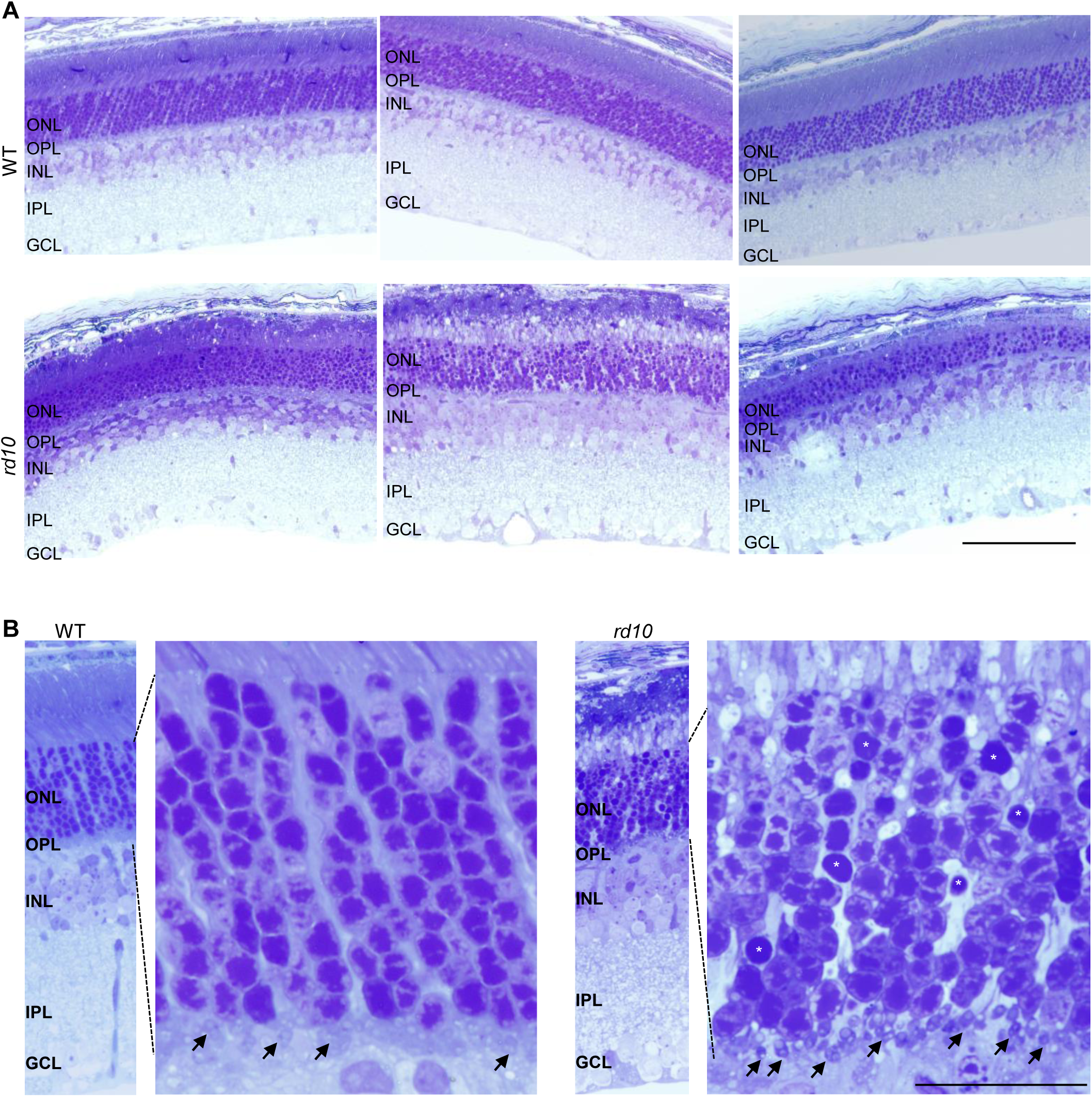
Representative semi-thin retinal sections. **A.** Semi-thin retinal sections collected at P21 from WT and *rd10* mice were prepared for electron microscopy analysis (Fig. 4D). Scale bar: 50 μm. **B.** Magnification of the indicated region. Arrows indicate photoreceptor presynaptic terminals. Asterisks indicate degenerating photoreceptors, as evidenced by the condensed nucleus. ONL, outer nuclear layer; OPL, outer plexiform layer; IPL, inner plexiform layer; INL, inner nuclear layer; NFL, nerve fiber layer. Scale bar: 100 μm.

**Supplementary Figure 6.**
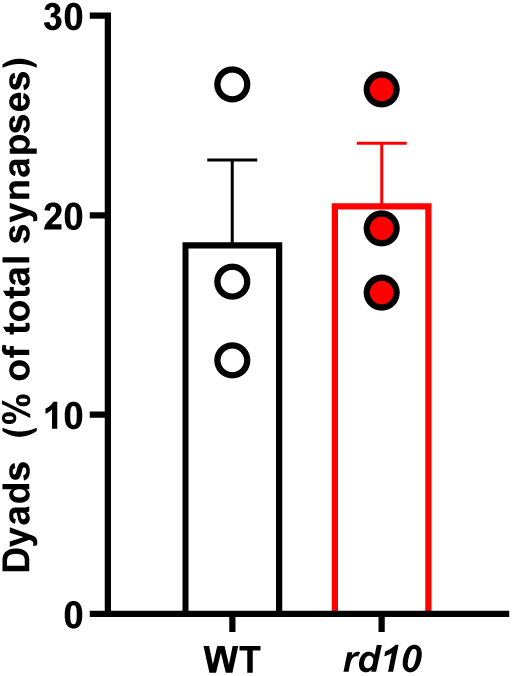
Quantification of the proportion of dyads. Data are presented as the mean + SEM. n=3 mice, ≥60 synapses per mouse.

**Supplementary Figure 7.**
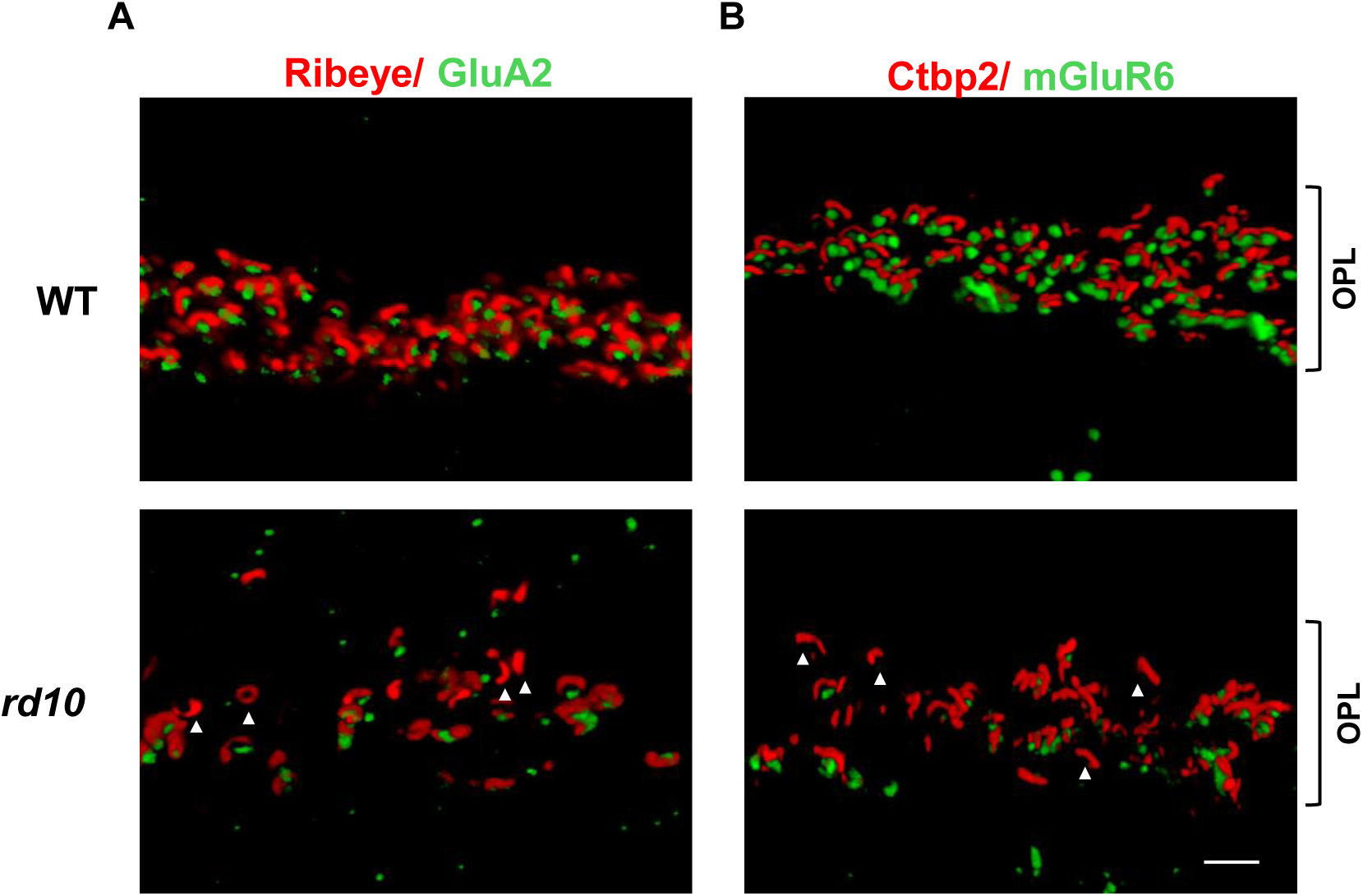
Representative 3D reconstructions of P23 WT and *rd10* synapses in the OPL. **A, B.** Retinal sections were co-immunostained for Ribeye (ribbon at presynaptic terminal, red) and the glutamate receptor subunit GluA2 (horizontal postsynaptic terminal, green) (A), or for Ctbp2 (ribbon at presynaptic terminal, red) and the glutamate receptor mGluR6 (bipolar postsynaptic terminal, green) (B). Arrowheads indicate ribbons (rod presynaptic terminals) without a post-synaptic partner. Scale bar: 6 μm.

**Supplementary Figure 8.**
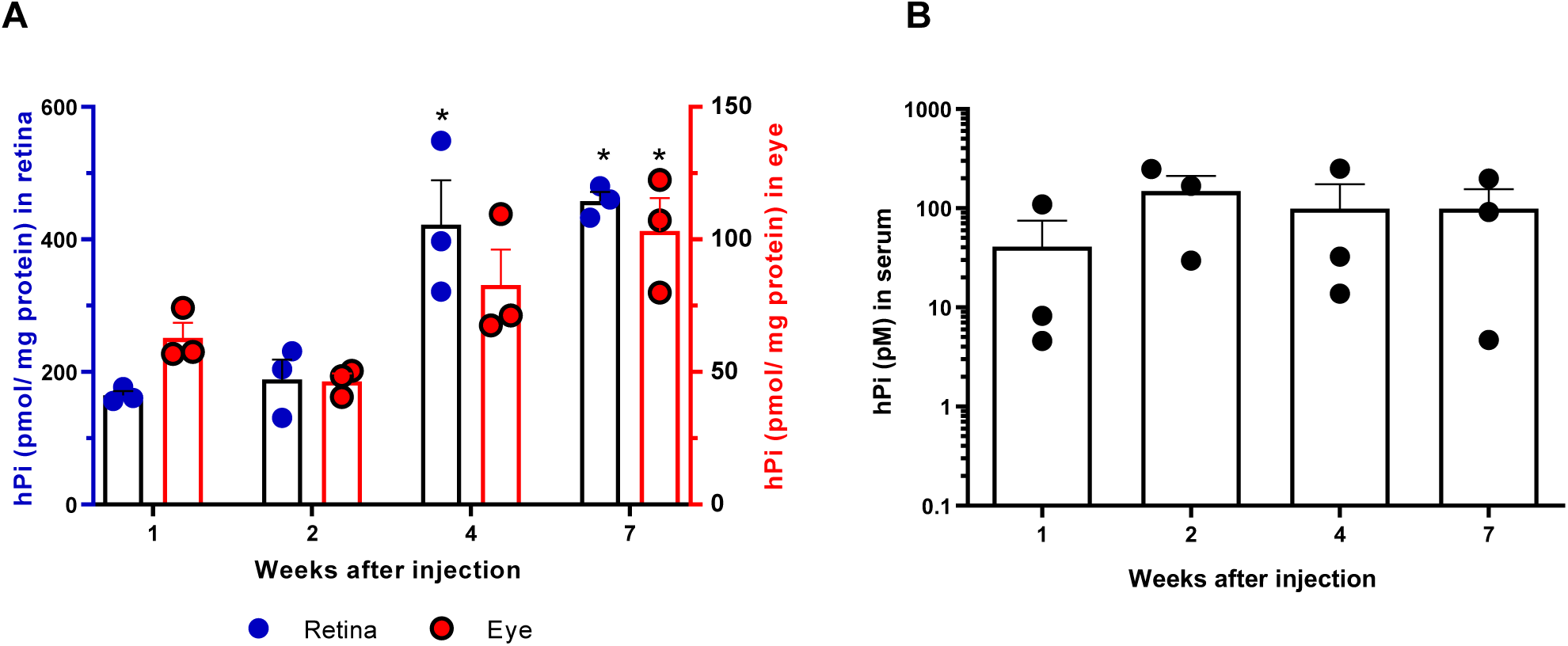
Determination of hPi levels after AAV-hPi treatment. *rd10* mice received a single intramuscular injection of AAV-hPi or AAV-null at P10. **A, B**. ELISA was performed to measure hPi in retinal and eye extracts (A) and in serum (B) at the indicated times. Data are presented as the mean + SEM. n=3 mice. \**p* ≤ 0.05 (1-way ANOVA with Dunnett’s multiple comparison test).

## ACKNOWLEDGEMENTS

The authors thank Cayetana Murillo, María D. Hernández-Fuentes, the staff at the CIB animal facility and CIB and UCM microscopy facilities for technical support; Nuria Forns and Ryan Steel for setting the conditions for the optomotor test and for synapse immunostaining; Noemi Alvarez Lindo for designing the diagrams; and José A. Esteban for supplying the anti-Glu2A antibody. This work was supported by grants from the Spanish MINECO (SAF2013-41059-R and SAF2016-75681-R to EJdlR, PI13-02098 to PdlV), the Instituto de Salud Carlos III and co-financed by the European Development Regional Fund ‘‘*A way to make Europe”/”Investing in your future*’’ (ERDF/ESF) (PI17/01601 to IL and PI18/01536 to CL), the Instituto de Salud Carlos III RETICS RD16/0019/0009 and the Madrid Regional Government B2017/BMD-3688 to IL. ASC is recipient of a UCM predoctoral fellowship (CT45/15).

## AUTHOR CONTRIBUTION

ASC, AHP, CI and MM performed experimental procedures. CL carried out and analyzed the electron microscopy studies. PdlV designed the functional studies and analyzed the data. FB designed and generated the AAV vectors. EJdlR and CHS designed and supervised the biological experiments. EJdlR and CHS analyzed and discussed the results. ASC, CL, CI, IL, PdlV, FB, EJdlR and CHS contributed to the writing and editing of the manuscript. All authors read and approved the final manuscript.

## THE PAPER EXPLAINED

### Problem

Retinitis pigmentosa is a hereditary retinal neurodegenerative condition that accounts for the majority of cases of congenital blindness. It is considered a rare disease, with a worldwide incidence of 1 per 3500 people. Although gene therapy interventions are considered the definitive therapeutic strategy, their application is hindered by the wide range of genes and mutations implicated in RP. Therefore, there is an urgent social and medical need to develop effective treatments that are independent of the causative mutation.

### Results

The insulin receptor is essential for the control of metabolism and glucose homeostasis. We show that the insulin receptor is predominantly expressed in the axons of retinal horizontal neurons. Moreover, the *rd10* mouse model of RP shows deficient insulin receptor signaling in horizontal neurons and aberrant synaptic contacts between rod photoreceptors and horizontal and bipolar cells. A gene therapy strategy to produce sustained systemic levels of proinsulin restored retinal insulin receptor signaling in *rd10* mice without affecting peripheral metabolism. Importantly, proinsulin treatment preserved photoreceptor synaptic connectivity and prolonged visual function.

### Impact

This study underscores the role of insulin receptor signaling in retinal dystrophies. Our results support gene therapy as a feasible therapeutic strategy to induce sustained proinsulin production. More importantly, they provide proof of concept of the therapeutic potential of proinsulin for the treatment of RP.

